# R-spondin 1 restores hypothalamic glucose-sensing and systemic glucose homeostasis via Wnt signaling in diet-induced obese mice

**DOI:** 10.64898/2026.03.26.714630

**Authors:** Ming-Liang Lee, Shawn He, Takashi Abe, Ching-Pu Chang, Ryosuke Enoki, Chitoku Toda

## Abstract

High-fat diet (HFD) feeding disrupts systemic glucose metabolism, yet the underlying neural mechanisms remain incompletely understood. Here, we demonstrate that glucose-excited (GE) neurons in the ventromedial hypothalamus (VMH^GE^) are essential for acute glucose regulation and that their function is compromised by HFD via structural synaptic remodeling. We found that HFD feeding suppresses canonical Wnt signaling and downregulates R-spondin 1 (RSPO1), a Wnt enhancer, in the VMH. This Wnt inhibition leads to a loss of dendritic spines and blunted glucose-sensing in VMH^GE^ neurons. Conversely, central administration of RSPO1 restores Wnt/β-catenin signaling, promotes synaptogenesis, and recovers neuronal glucose responsiveness. Consequently, RSPO1 treatment ameliorates HFD-induced glucose intolerance by enhancing peripheral glucose utilization. These findings identify the RSPO1-Wnt signaling axis as a critical regulator of VMH neuronal plasticity and metabolic homeostasis, providing a mechanistic link between diet-induced synaptic pathology and systemic metabolic dysfunction.

**Highlights:**

- Glucose-excited neurons in VMH were labeled with TRAP
- VMH glucose-excited neurons regulates systemic glucose metabolism
- Wnt signaling regulates synaptogenesis in VMH and maintain neuronal glucose-sensitivity
- R-spondin1 recovers VMH neuronal glucose sensitivity in HFD fed obese mice

## Introduction

Precise regulation of blood glucose is essential for survival, particularly given that the brain accounts for roughly 25% of total body glucose consumption^1^. The ventromedial hypothalamus (VMH) serves as a primary effector site in this regulation, modulating the autonomic nervous system to adjust hepatic glucose production and glucose uptake in muscles^2,3^. Importantly, the VMH appears to exert opposing effects on glucose metabolism depending on the specific subpopulation of neurons targeted or the activation method used. For instance, optogenetic activation of the steroidogenic factor 1 (SF1) neuronal population or estrogen receptor alpha (ERα) has been shown to increase blood glucose levels by altering counter hormones, such as glucagon, catecholamines, and corticosterone^4,5^. In contrast, selective excitation of the dorsomedial subregion (dmVMH) using Designer Receptors Exclusively Activated by Designer Drugs (DREADDs) promotes systemic glucose utilization while maintaining stable blood glucose levels^6^. These divergent metabolic outcomes highlight the cellular and regional complexity within the VMH, suggesting that distinct neural circuits are responsible for fine-tuning glucose homeostasis.

Hypothalamic glucose-sensing neurons are believed to detect blood glucose levels and modulate systemic glucose metabolism to maintain energy homeostasis^7^. Within the VMH, two distinct populations exist: glucose-excited (GE) and glucose-inhibited (GI) neurons, which are excited or inhibited by the increase in glucose levels, respectively^8^. The coexistence of these heterogeneous populations with opposing functions presents a challenge for physiological analysis. While the role of GI neurons in the recovery from hypoglycemia is well-documented^7,8^, the specific contribution of GE neurons remains less understood. We previously demonstrated that inhibition of mitochondrial uncoupling protein 2 (UCP2)-mediated GE neuronal activity in the VMH decreases glucose tolerance^9^; however, the lack of appropriate biomarkers has restricted further investigation into these neurons.

Wnt signaling is a critical regulator of tissue maintenance and remodeling^10^. In humans, variants of *TCF7L2*, a key component of the Wnt pathway, are associated with an increased risk of type 2 diabetes, suggesting a link between Wnt signaling and glucose metabolism^11,12^. In the brain, Wnt signaling regulates the morphogenesis of axons, dendrites, and synapses, thereby influencing neuronal function^13–15^. Moreover, Wnt signaling has been shown to enhance neuronal glucose metabolism, protecting against synaptic loss^16^. Here, we established a method to label GE neurons and demonstrated that their role in regulating systemic glucose metabolism is controlled by Wnt signaling. Notably, high-fat diet feeding impairs GE neuron function through this mechanism.

## Results

### Glucose-excited neurons in the VMH were conditionally labeled by TRAP

To specifically investigate GE neurons in the VMH, we utilized the Targeted Recombination in Active Populations (TRAP) method to conditionally label these neurons *in vivo* (Fig. 1a)^17^. Briefly, glucose (3 g/kg) was administered intraperitoneally (i.p.) four times at 20-min intervals into Arc-CreER::Ai14 bi-transgenic mice to activate GE neurons (SFig1a). The activity-dependent *Arc* promoter drives CreER expression in activated neurons; subsequently, 4-hydroxytamoxifen (4-OHT) was i.p. injected to induce the nuclear translocation of CreER and trigger tdTomato expression (SFig. 1a).

**Figure 1.**
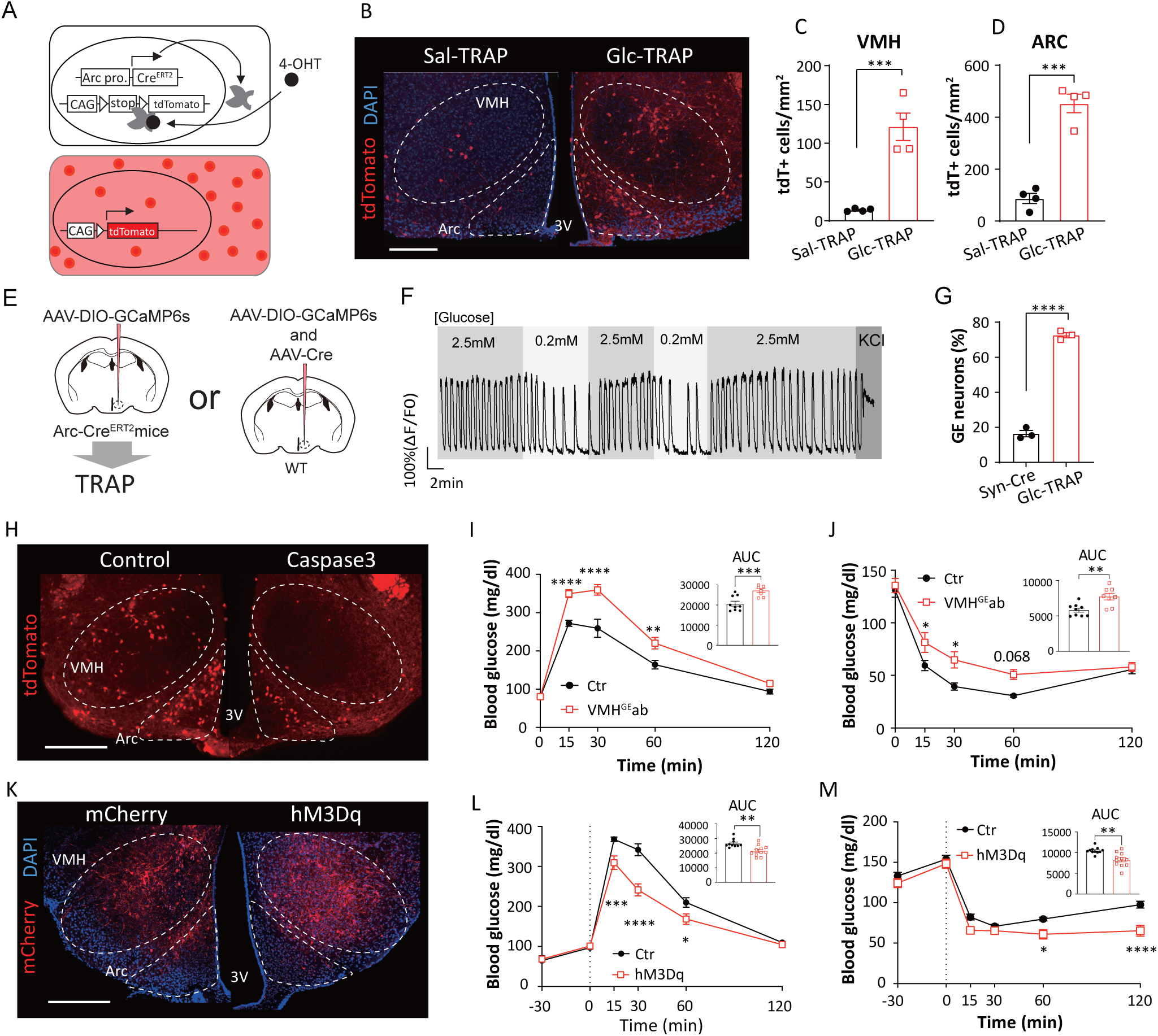
Labeling of glucose-excited (GE) neurons in the VMH. (A) Schematic illustration of the Targeted Recombination in Active Populations (TRAP) method used to label GE neurons. Briefly, the expression of CreER is driven by the activity-regulated cytoskeleton-associated protein (Arc) promoter in response to glucose stimulation. 4-OHT injection induces the nuclear translocation of CreER, which triggers the expression of tdTomato. (B) Representative micrographs of TRAP-labeled neurons in the hypothalamus following glucose or saline injection. Scale bar, 200 μm. (C and D) Number of TRAP-labeled neurons in the VMH (C) and ARC (D) (n = 4 per group). (E-G) Functional characterization of Glc-TRAPed neurons via slice calcium imaging. (E) Schematic strategy for calcium imaging of Glc-TRAPed neurons. (F) Representative calcium traces of a Glc-TRAPed neuron in response to changes in extracellular glucose concentration. (G) Proportion of glucose-excited (GE) neurons within the Glc-TRAPed population in the VMH (n = 3 mice). (H–J) Impact of VMH^GE^ ablation on systemic glucose metabolism. (H) Validation of VMH^GE^ ablation. Representative micrograph showing the reduction of TRAP-labeled neurons in the VMH following caspase-3-mediated ablation. Scale bar, 200 μm. (I and J) Intraperitoneal glucose tolerance test (GTT) (I) and insulin tolerance test (ITT) (J) in control and VMH^GE^ -ablated (VMH^GE^ab) mice (VMH^GE^ab: n=8, Control: n=9). (K–M) Impact of chemogenetic activation of VMH^GE^ neurons on glucose metabolism. (K) Representative micrograph of the hypothalamus showing TRAP-mediated expression of hM3Dq-mCherry. Scale bar, 200 μm. (L and M) Glucose tolerance test (L) and insulin tolerance test (M) following clozapine injection in control and hM3Dq-expressing mice (hM3Dq: n=11, Control: n=10). All data represent the mean ± SEM; * *p*<0.05; ** p<0.01; *** p<0.001; **** p<0.0001.

A substantial number of neurons in both the VMH and the arcuate nucleus (ARC) were “TRAPed” (expressing tdTomato) following glucose and 4-OHT injection (Glc-TRAP). In contrast, only a few TRAPed neurons were observed in control mice injected with saline and 4-OHT (Sal-TRAP) (Fig. 1B-1D). To determine whether the TRAPed neurons respond to glucose stimulation, mice were sacrificed following an i.p. glucose injection, and brain sections were immunostained for glucose-induced c-fos expression (SFig1 B-D). In Sal-TRAP mice, less than 20% of TRAPed neurons in VMH were c-fos positive, whereas this ratio was significantly higher in Glc-TRAP mice, exceeding 50% (SFig. 1C). However, the proportion of c-fos positive TRAPed neurons in the ARC was not significantly different between Glc-TRAP and Sal-TRAP mice (Fig. 1D). Systemic glucose injection can induce the secretion of hormones, such as insulin, which may also modulate neuronal activity. To eliminate potential confounding effects from circulating hormones, we performed slice calcium imaging to confirm whether the TRAPed neurons in the VMH are indeed GE neurons (Fig. 1E to 1G). We injected AAV-DIO-GCaMP6s into the VMH to selectively express the calcium indicator in TRAPed neurons. Control mice were co-injected with AAV-hSyn-Cre and AAV-DIO-GCaMP6s to express GCaMP6s non-specifically in VMH neurons (Fig. 1E). Neuronal activity under different glucose concentrations was recorded via slice calcium imaging. The activity of TRAPed neurons decreased when the perfusion solution was switched from 2.5 mM to 0.2 mM glucose and recovered upon reintroduction of 2.5 mM glucose, confirming that these TRAPed neurons are GE neurons (Fig. 1F). In control mice, approximately 15% of non-specific VMH neurons were identified as GE neurons. In contrast, over 70% of Glc-TRAPed neurons in the VMH displayed GE characteristics (Fig. 1G). Therefore, TRAP is a practical strategy for labeling GE neurons in the VMH *in vivo*.

### TRAPed GE neurons in the VMH regulate systemic glucose metabolism

The VMH plays a pivotal role in regulating systemic glucose metabolism; however, the specific contribution of GE neurons in the VMH (VMH^GE^) to this control remains unclear. To investigate the role of VMH^GE^ neurons, we assessed glucose metabolism following the specific ablation of these neurons (Fig. 1H–1J). We injected AAV-DIO-taCasp3 into the VMH to selectively ablate VMH^GE^ neurons via TRAP-induced overexpression of active caspase-3. The number of tdTomato-positive neurons was significantly reduced in the VMH, but not in the ARC, following the overexpression of active caspase-3, confirming that the TRAPed neurons in the VMH, rather than those in the ARC, were selectively ablated (Fig. 1H). Mice with VMH^GE^ ablation exhibited glucose intolerance and insulin resistance compared to control mice (Fig. 1I and 1J). However, body weight and fasting blood glucose remained unchanged (SFig. 2A to 2C). These results suggest that VMH^GE^ neurons regulate acute glucose metabolism rather than long-term energy homeostasis.

Next, we expressed hM3Dq in VMH^GE^ neurons to selectively activate them using DREADD-based chemogenetics (Fig. 1K to 1M). DREADD-mediated activation of VMH^GE^ neurons improved glucose tolerance and insulin sensitivity (Fig. 1L and 1M). Consistent with the ablation results, basal blood glucose levels were not affected by VMH^GE^ activation (SFig. 2D and 2E). Collectively, these findings indicate that VMH^GE^ neurons are necessary and sufficient to regulate systemic glucose metabolism without altering basal blood glucose levels.

### High-fat diet feeding impairs systemic glucose metabolism by blunting the glucose-sensing of GE neurons in the VMH

To understand the relationship between systemic glucose metabolism and the functions of VMH^GE^ neurons under high-fat diet (HFD) conditions, VMH^GE^ -ablated mice were fed a HFD for 3 months. This duration is sufficient to disturb systemic energy balance, resulting in elevated blood glucose levels in both fed and fasted states (Fig. 2A and 2B). Consistent with above results, ablation of VMH^GE^ neurons did not affect basal blood glucose levels, even after HFD feeding, further confirming that these neurons are not involved in long-term glucose homeostasis (Fig. 2A and 2B).

**Figure 2.**
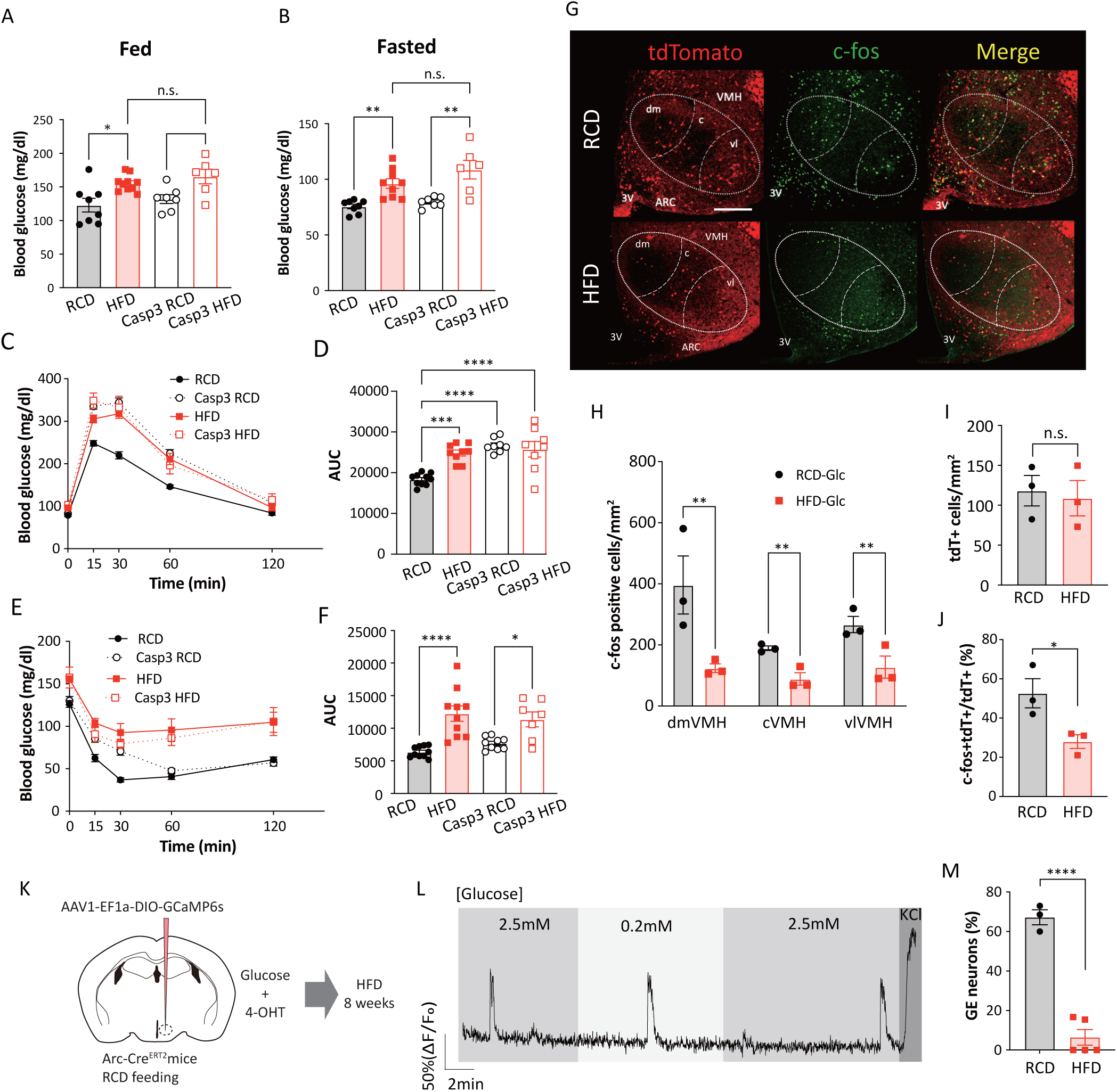
High-fat diet impairs glucose sensing of VMH^GE^. (A and B) Basal blood glucose levels in control and VMH^GE^-ablated mice fed RCD or HFD for 2 months. (A) *Ad libitum* fed state and (B) fasted state. (RCD: n=8, HFD: n=10, Casp3 RCD: n=7, Casp3 HFD: n=6). (C-F) Systemic glucose metabolism in VMH^GE^-ablated mice fed RCD or HFD. (C) Time course of blood glucose levels and (D) area under the curve (AUC) during GTT (RCD: n=10, HFD: n=10, Casp3 RCD: n=8, Casp3 HCD: n=8). (E) Time course of blood glucose levels and (F) AUC during ITT (RCD: n=10, HFD: n=10, Casp3 RCD: n=8, Casp3 HCD: n=8). (G-J) Glucose induced c-fos expression in RCD- or HFD-fed mice. (G) Representative micrographs of c-fos immunohistochemistry and tdTomato expression in the dorsomedial (dm), central (c), and ventrolateral (vl) subregions of the VMH and arcuate nucleus (ARC) following glucose injection. Scale bar, 200 μm. (H) Quantification of c-fos positive cells in subregions of the VMH. (I) Number of tdTomato-positive (tdT+) neurons in the VMH. (J) Proportion of c-fos/tdTomato double-positive cells in the VMH. (K–M) Assessment of VMH^GE^ glucose sensitivity by slice calcium imaging. (K) Schematic of the experimental preparation. (L) Representative calcium traces of a Glc-TRAPed neuron from an HFD-fed mouse in response to changes in extracellular glucose concentration. (M) Proportion of GE neurons among tdTomato-positive neurons in the VMH (RCD: n=3, HFD: n=5). All data represent the mean ± SEM; * *p*<0.05; ** p<0.01; *** p<0.001; **** p<0.0001.

In control mice, three months of HFD feeding impaired both glucose tolerance and insulin sensitivity (Fig. 2C to 2F). Strikingly, HFD feeding did not exacerbate glucose intolerance in VMH^GE^-ablated mice; the glucose tolerance of HFD-fed VMH^GE^-ablated mice was comparable to that of RCD-fed VMH^GE^-ablated mice and HFD-fed control mice (Fig. 2C and 2D). This lack of additive effect suggests that HFD-induced glucose intolerance is mediated primarily by the impairment of VMH^GE^ neurons. In contrast, insulin sensitivity in VMH^GE^-ablated mice was significantly worsened by HFD feeding (Fig. 2E and 2F), indicating that HFD affects insulin sensitivity via mechanisms independent of VMH^GE^ neurons. Collectively, these results suggest that VMH^GE^ neurons specifically regulate glucose metabolism under hyperglycemic condition.

We next investigated how HFD affects neuronal glucose-sensing in the VMH (Figs. 2G to 2J). The VMH^GE^ were TRAPed followed by 3 months of HFD feeding, and were sacrificed after glucose injection. The number of glucose-induced c-fos positive neurons was significantly lower in the dorsomedial (dm)- and ventrolateral (vl)-VMH after HFD feeding, indicating impaired glucose responsiveness of the VMH (Figs. 2G and 2H). Furthermore, the ratio of c-fos-positive TRAPed neurons was decreased while the population of TRAPed neurons remained unchanged, which suggests that HFD blunts the glucose responsiveness of VMH^GE^ rather than causing a loss of these neurons (Figs. 2I and 2J). Feeding HFD for 3 months did not change the population of TRAPed neurons or glucose-induced c-fos expression in the ARC (SFig. 3).

To determine if HFD impaired the glucose responsiveness of VMH^GE^, we conducted slice calcium imaging on VMH^GE^ after HFD feeding (Figs. 2K to 2M). VMH^GE^ neurons largely failed to respond to glucose after HFD feeding, suggesting that HFD impairs their glucose-sensing function (Fig. 2L). The ratio of GE neurons in the VMH among TRAPed neurons decreased dramatically from ∼70% in RCD-fed mice to only ∼10% after HFD feeding (Fig. 2M).

### HFD deteriorates glucose-sensing and spinogenesis of VMH^GE^ with inhibited wnt signaling

To elucidate the molecular mechanisms underlying HFD-induced impairment of glucose sensitivity in VMH^GE^ neurons, we performed RNA-seq on TRAPed VMH^GE^ neurons from mice fed an RCD or HFD (Fig. 3A). Gene Ontology (GO) analysis was conducted to uncover the major molecular changes. Gene sets related to the postsynapse and mitochondria were markedly affected by HFD feeding (SFig. 4A and Fig. 3B). mRNA expression associated with synaptic structures was upregulated, while mitochondrial genes were downregulated by HFD feeding (SFig. 4A). Among them, genes associated with post-synapses, especially asymmetric (excitatory) synapses, were the most significantly enriched (Fig. 4B). Unexpectedly, despite higher *Dlg4* (the gene encoding PSD95) mRNA expression (SFig. 4B), the density of dendritic spines—the primary sites of asymmetric synapses—was lower in HFD-fed mice compared to RCD mice, (Figs. 3C and 3D). To reconcile the discrepancy between dendritic spine density and mRNA expression, we quantified PSD95 protein in the mediobasal hypothalamus by western blotting. Consistent with the decreased dendritic spines, PSD95 protein levels were lower in HFD-fed mice (Fig. 3E). This suggests that HFD feeding impairs the translation of mRNA, a phenomenon that parallels the HFD-induced translational suppression observed in pancreatic β-cells^18,19^. The gene set for ribosome subunits was downregulated by HFD feeding, which likely disturbed ribosome biogenesis and resulted in translational impairment (SFig. 4A).

**Figure 3.**
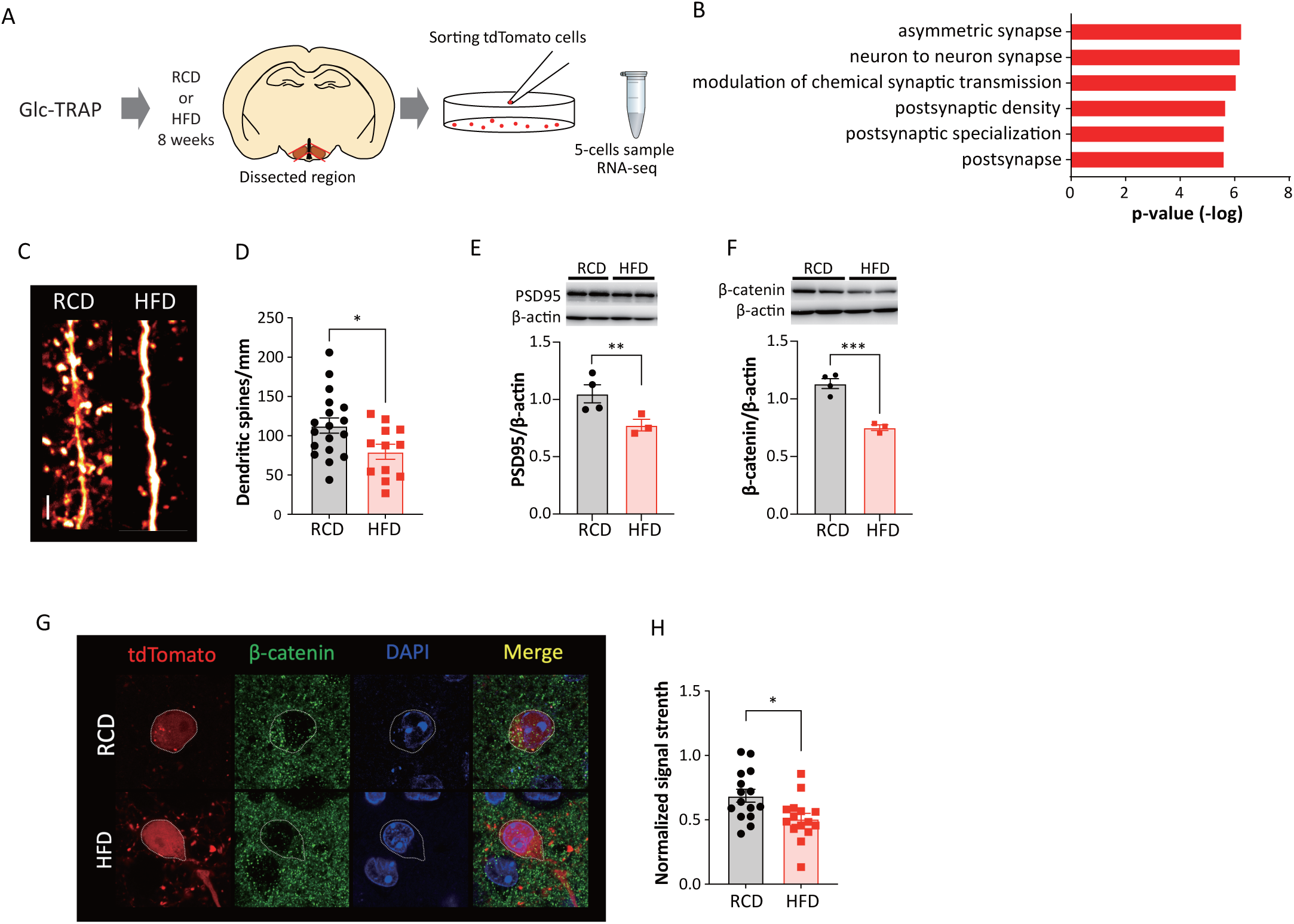
High-fat-diet feeding disturbs synaptogenesis and blocks Wnt signaling. (A) Schematic illustration of the RNA-seq workflow for VMH^GE^ neurons isolated from RCD- or HFD-fed mice. (B) Top 5 significantly enriched Gene Ontology (GO) terms identified by the analysis. (C and D) Analysis of dendritic spine density in VMH^GE^ neurons. (C) Representative micrographs of dendritic shafts and spines in VMH^GE^ neurons. Scale bar, 5μm. (D) Quantification of dendritic spine density. (E and F) Western blot analysis of PSD95 (E) and β-catenin (F) protein levels in the mediobasal hypothalamus (MBH). (G and H) Evaluation of intracellular β-catenin levels in VMH^GE^ neurons. (G) Representative immunohistochemistry images showing β-catenin expression in VMH^GE^ neurons. (H) Quantification of β-catenin signal intensity in VMH^GE^ neurons (RCD: n = n=18 from 3 mice, HFD: n = 12 from 3 mice). All data represent the mean ± SEM; * *p*<0.05; ** *p*<0.01; *** *p*<0.001; **** *p*<0.0001.

Wnt signaling is a key factor regulating dendritic spinogenesis and can be inhibited by HFD in the hippocampus^20,21^. We observed that HFD feeding significantly reduced the protein levels of β-catenin, a central component of canonical Wnt signaling, in the mediobasal hypothalamus (Fig. 3F), suggesting an impairment of this pathway in the hypothalamus. Strikingly, despite the upregulation of *Ctnnb1* mRNA (encoding β-catenin) (SFig. 4D)—a pattern reminiscent of the synaptic gene discrepancy—intracellular β-catenin signaling was reduced in VMH^GE^ neurons (Fig. 3G and 3H).

Therefore, we next investigated whether Wnt signaling is required for VMH^GE^ neuronal function. To test this, we administered the Wnt inhibitor Dkk1 intracerebroventricularly (i.c.v.) to lean mice. I.c.v. injection of Dkk1 decreased β-catenin protein in the VMH^GE^ neurons (Figs. 4A and 4B), confirming that Dkk1 inhibits Wnt signaling in these neurons. Glucose-induced c-fos expression was decreased in Dkk1-injected mice, mainly in the dorsomedial VMH (dmVMH) subregion (Figs. 4C and 4D). Dkk1 also impaired glucose-induced c-fos expression in VMH^GE^ neurons, indicating that inhibition of Wnt signaling blunts the glucose-sensing of these neurons in dmVMH (Fig. 4E).

**Figure 4.**
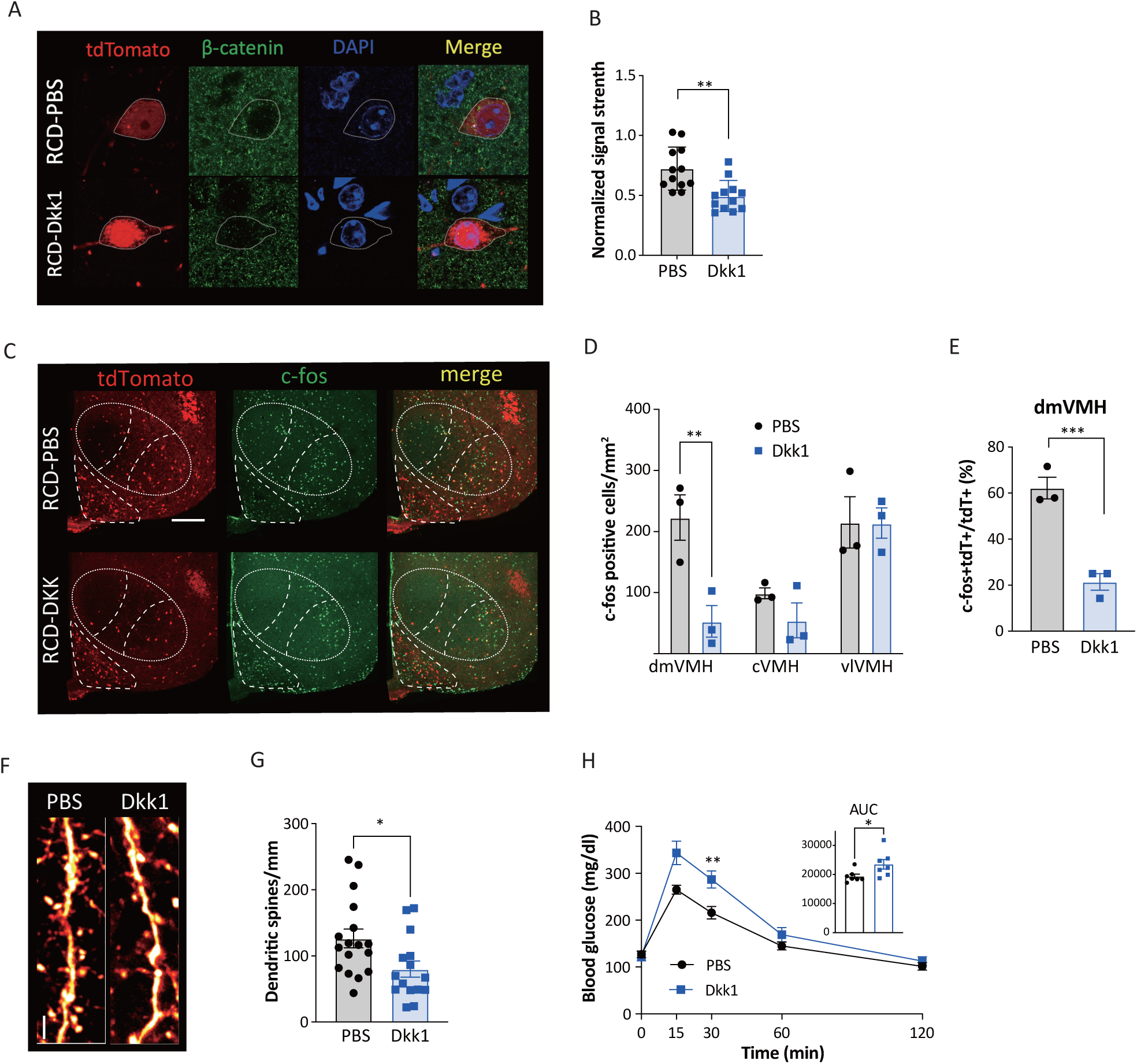
Blocking Wnt signaling impairs glucose sensing of VMH^GE^ in lean mice. (A and B) Impact of central administration of the Wnt inhibitor Dkk1 on β-catenin levels in VMH^GE^ neurons. (A) Representative immunohistochemistry images showing β-catenin expression in VMH^GE^ neurons following intracerebroventricular (i.c.v.) injection of Dkk1 or PBS. (B) Quantification of β-catenin signal intensity in VMH^GE^ neurons. (C–E) Glucose-induced c-fos expression in the VMH following Dkk1 administration. (C) Representative micrographs of c-fos expression after i.p. glucose injection in mice pre-treated with i.c.v. Dkk1. Scale bar, 200μm. (D) Quantification of c-fos-positive cells in each VMH subregion. (E) Proportion of c-fos/tdTomato double-positive cells in the dmVMH. (F and G) Effect of i.c.v. Dkk1 injection on dendritic spine density in VMH^GE^ neurons. (F) Representative micrographs of dendritic shafts and spines. Scale bar, 5μm. (G) Quantification of dendritic spine density. PBS: n = 18 cells from 3 mice; Dkk1: n = 15 cells from 3 mice). (H) Glucose tolerance test following i.c.v. injection of Dkk1 (PBS: n = 7, Dkk1: n = 7). All data represent the mean ± SEM; * *p*<0.05; ** *p*<0.01; *** *p*<0.001.

Inhibition of Wnt signaling by Dkk1 has been shown to induce the withdrawal of dendritic spines in the hippocampus^22^. Similarly, the density of dendritic spines decreased in VMH^GE^ neurons after Dkk1 injection (Figs. 4F and 4G). Furthermore, central Dkk1 administration impaired glucose tolerance, demonstrating that inhibiting Wnt signaling is sufficient to disrupt systemic glucose metabolism (Fig. 4H). Therefore, Wnt signaling in VMH^GE^ neurons, which is inhibited by HFD, plays a crucial role in regulating synaptogenesis, glucose-sensing, and peripheral glucose metabolism.

### R-spondin 1 restores glucose sensing and increases spinogenesis of VMH^GE^ neurons in HFD-fed mice

R-spondin 1 (RSPO1), which is highly expressed in the VMH, is a potent enhancer of the canonical Wnt pathway^23^. RSPO1 stabilizes Wnt signaling by preventing the internalization and degradation of Frizzled receptors^24,25^. In the brain, the receptor for RSPO1, Leucine-rich repeat-containing G protein-coupled receptor 4 (LGR4), exhibits restricted expression in regions such as the amygdala, paraventricular thalamus (PVT), habenula, and hypothalamus^23^. Within the hypothalamus, LGR4 expression is localized to the VMH, ARC, median eminence, and ependymocytes^23^. RSPO1-expressing neurons in the VMH suppress food intake by activating CART neurons in the ARC^26^. We found that HFD feeding led to a suppression of RSPO1 mRNA in the VMH (Supplementary Fig. 4D). This change is consistent with decreased β-catenin levels, supporting the idea that HFD inhibits Wnt signaling.

To determine whether central RSPO1 affects the glucose responsiveness of the VMH, we administered RSPO1 i.c.v. followed by an i.p. injection of glucose. Mirroring the inhibitory effects of central Dkk1on dmVMH, i.c.v. injection of RSPO1 increased glucose-induced c-fos expression in the dmVMH, but not in the central (cVMH) or ventrolateral VMH (vlVMH) (Fig. 5A, 5B, and 4D). The increased proportion of c-fos-positive VMH^GE^ neurons demonstrates that RSPO1 improves the glucose responsiveness of this population (Fig. 5C). Glucose-induced c-fos expression was unaffected by RSPO1 in other LGR4-expressing regions, such as the ARC, amygdala, and paraventricular thalamus (Supplementary Fig. 5). Thus, RSPO1selectively increases glucose-induced neuronal activity in the VMH. Furthermore, RSPO1 treatment increased dendritic spine density in VMH^GE^ neurons of HFD-fed mice (Fig. 5D and 5E). To determine whether RSPO1 restores glucose sensing in VMH^GE^ neurons, we performed slice calcium imaging on HFD-fed mice following RSPO1 injection (Fig. 5F and 5G). The number of glucose-responsive VMH^GE^ neurons was increased by RSPO1 injection, indicating that RSPO1 restores glucose-sensing function in these neurons (Fig. 5G).

**Figure 5.**
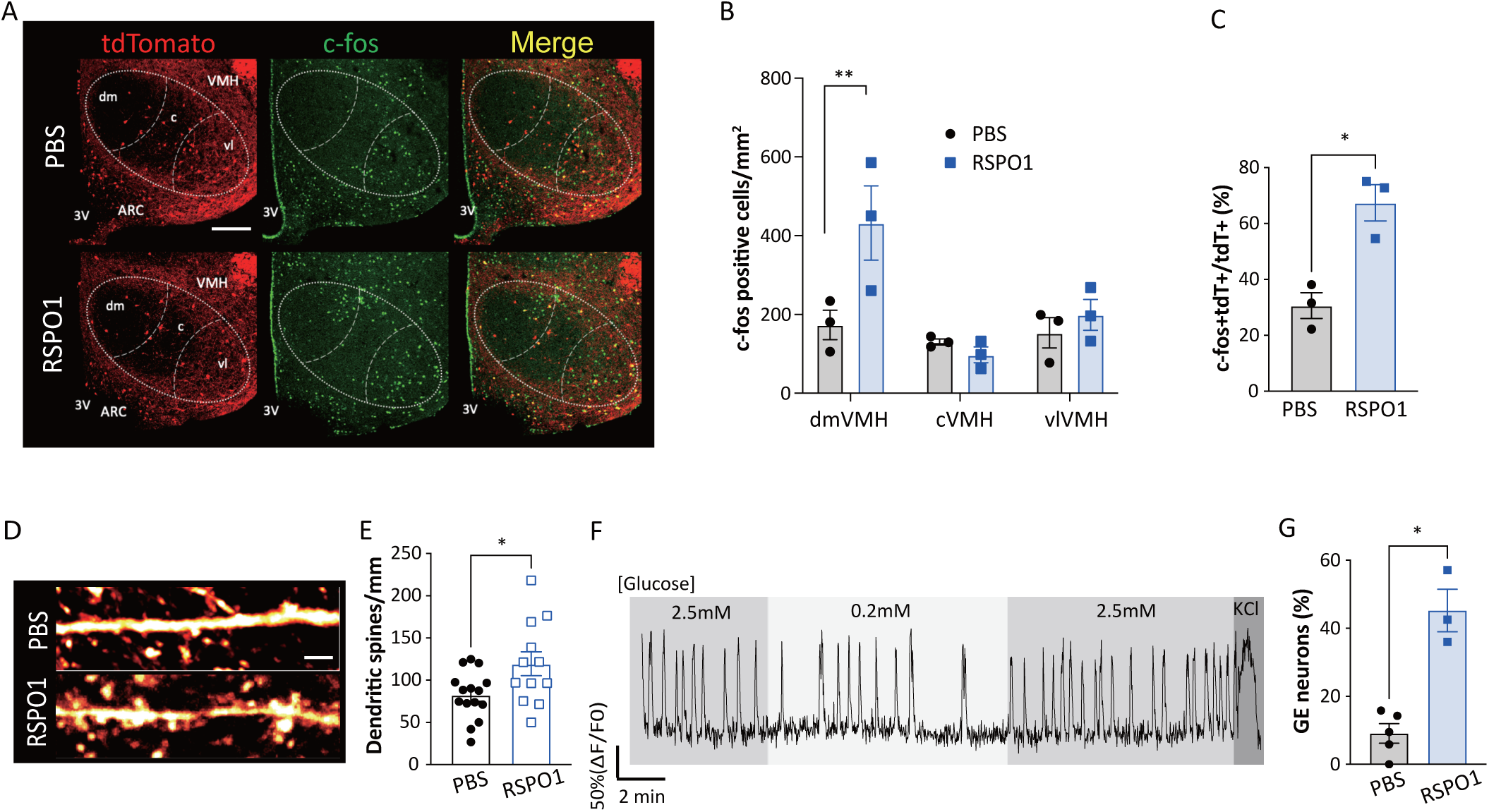
R-spondin1 recovers glucose sensing of VMH^GE^ in HFD fed mice. (A–C) Rescue of glucose-induced c-fos expression in HFD-fed mice by central RSPO1 administration. (A) Representative micrographs of c-fos expression in the VMH following i.p. glucose injection in HFD-fed mice treated with i.c.v. RSPO1. Scale bar, 200μm. (B) Quantification of c-Fos-positive cells in each VMH subregion. (C) Proportion of c-fos/tdTomato double-positive cells in the dmVMH. (D and E) Restoration of dendritic spine density in VMH^GE^ neurons by i.c.v. RSPO1. (D) Representative micrographs of dendritic shafts and spines in VMH^GE^ neurons from HFD-fed mice treated with RSPO1. Scale bar, 5μm. (E) Quantification of dendritic spine density (PBS: n = 15 cells from 3 mice; RSPO1: n = 12 cells from 3 mice). (F and G) Recovery of VMH^GE^ glucose sensitivity via RSPO1 treatment. (F) Representative calcium trace of a Glc-TRAPed neuron from an RSPO1-treated HFD-fed mouse in response to glucose. (G) Proportion of glucose-excited (GE) neurons among TRAP-labeled neurons in the VMH. All data represent the mean ± SEM; * *p*<0.05.

### Central RSPO1 administration improves peripheral glucose utilization

We next investigated whether the RSPO1-mediated improvement in VMH^GE^ glucose sensing is sufficient to improve whole-body glucose metabolism. Following i.c.v. injection of RSPO1, glucose tolerance and insulin sensitivity in HFD-fed mice were markedly improved (Fig. 6A and 6B). However, basal blood glucose levels in both ad libitum and fasted mice were unaffected by RSPO1 injection (Supplementary Fig. 6). We next conducted hyperinsulinemic-euglycemic clamp (Fig. 6C to 6P). HFD-fed mice injected with RSPO1 i.c.v. exhibited a higher glucose infusion rate (GIR) compared to PBS-injected controls (Fig. 6D and 6E). The rate of disappearance (Rd) and glycolysis were significantly higher in RSPO1-treated mice during the clamp period (Fig. 6F and 6G). Inhibition of endogenous glucose production (EGP) was not affected by RSPO1 (Fig. 6H and 6I). Therefore, RSPO1 improves the function of VMH^GE^ neurons to enhance glucose utilization rather than inhibiting EGP. Consistent with these results, 2DG uptake was increased in the soleus muscle, heart, and spleen (Fig. 6J to 6L). Interestingly, increased 2DG uptake in the hypothalamus suggests improved neuronal activity and function (Fig. 6M). The 2DG uptake in EWAT, BAT, and the cerebral cortex remained unchanged (Fig. 6N to 6P).

**Figure 6.**
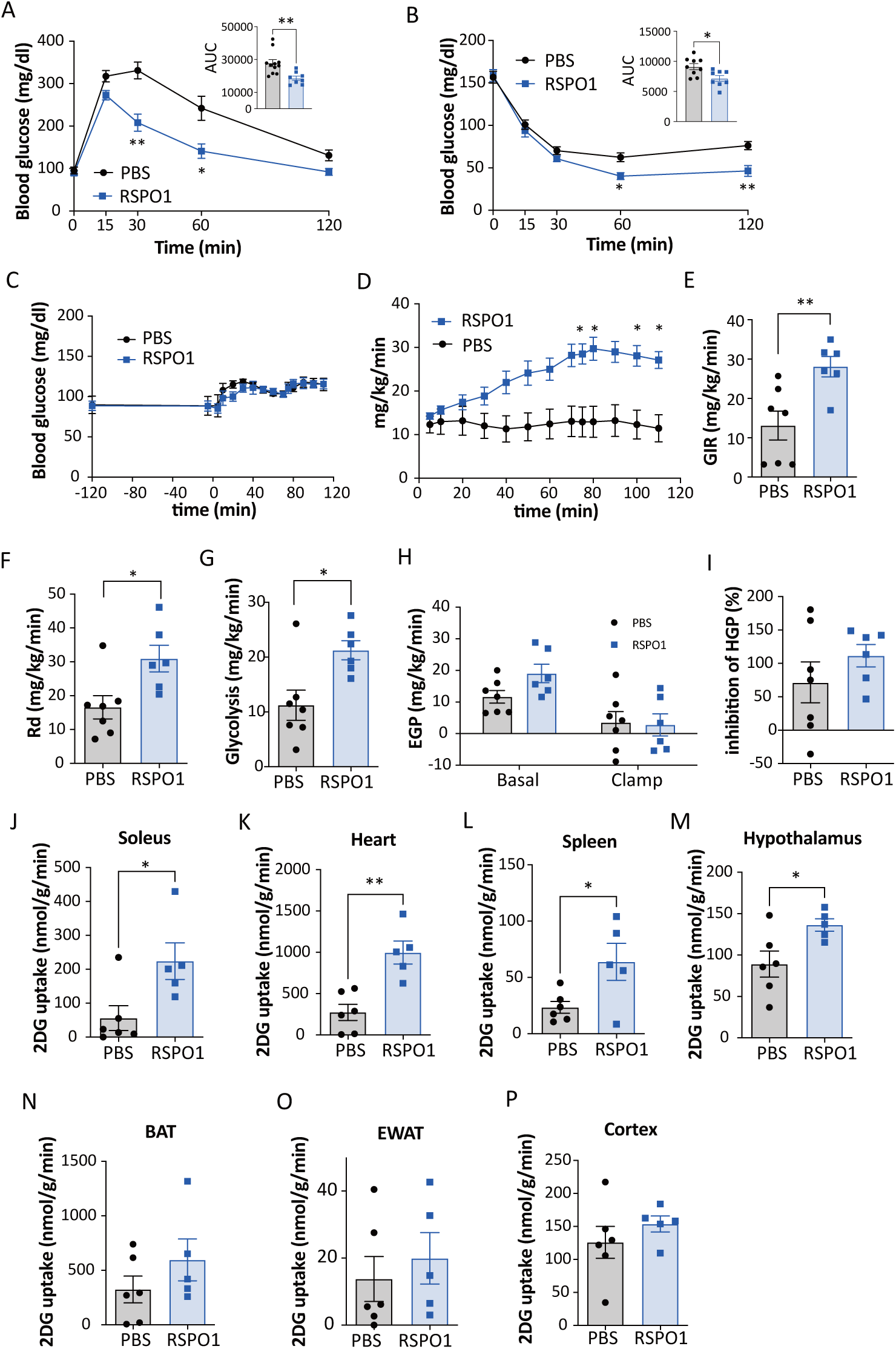
Central RSPO1 administration rescues HFD-induced impairment of glucose homeostasis. (A and B) Systemic glucose homeostasis in HFD-fed mice following central RSPO1 administration. (A) Glucose tolerance test and (B) insulin tolerance test following i.c.v. injection of RSPO1 (n=8) or PBS (n=8). (C–I) Hyperinsulinemic-euglycemic clamp analysis in HFD-fed mice treated with i.c.v. RSPO1 (n=6) or PBS (n=7). (C) Time course of blood glucose levels during the clamp. (D) Time course of the glucose infusion rate (GIR) required to maintain euglycemia. (E) Average GIR during the steady-state period (75–115 min). (F) Rate of whole-body glucose disappearance (Rd). (G) Rate of whole-body glycolysis. (H) Endogenous glucose production (EGP) under basal and clamp periods. (I) Percent suppression of EGP by insulin. (J–P) Tissue-specific glucose uptake determined by 2-[^14^C]-deoxy-D-glucose (2-DG) accumulation during the clamp period in soleus muscle (J), heart (K), spleen (L), hypothalamus (M), brown adipose tissue (BAT) (N), epididymal white adipose tissue (EWAT) (O), and cerebral cortex (P). All data represent the mean ± SEM; * *p*<0.05; ** *p*<0.01.

### R-spondin 1 regulates VMH^GE^ function by enhancing Wnt signaling

To confirm the involvement of Wnt signaling in the RSPO1-mediated regulation of systemic metabolism, we co-injected the Wnt antagonist Dkk1 in HFD-fed mice. This treatment abolished the RSPO1-induced improvement in glucose tolerance, reverting to levels comparable to control mice (Figs. 7A and 7B). The RSPO1-induced increase in c-fos expression in the dmVMH after glucose injection was also blocked by co-injection of Dkk1 (Figs. 7C and 7D). Similarly, the enhanced glucose-sensing in VMH^GE^ neurons by RSPO1 was abolished by Dkk1 co-injection (Fig. 7E). Furthermore, the effect of RSPO1 on increasing dendritic spine density was abrogated by Dkk1 (Figs. 7F and 7G). Finally, to directly confirm whether RSPO1 activates Wnt signaling in this context, we quantified β-catenin levels in VMH^GE^ neurons (Figs. 7H and 7I). RSPO1 injection increased the signal intensity of β-catenin, whereas co-injection of Dkk1 abolished this effect (Fig. 7H). Taken together, these results suggest that RSPO1 promotes synaptogenesis and restores glucose-sensing in VMH neurons by enhancing Wnt/β-catenin signaling, thereby improving systemic glucose metabolism.

**Figure 7.**
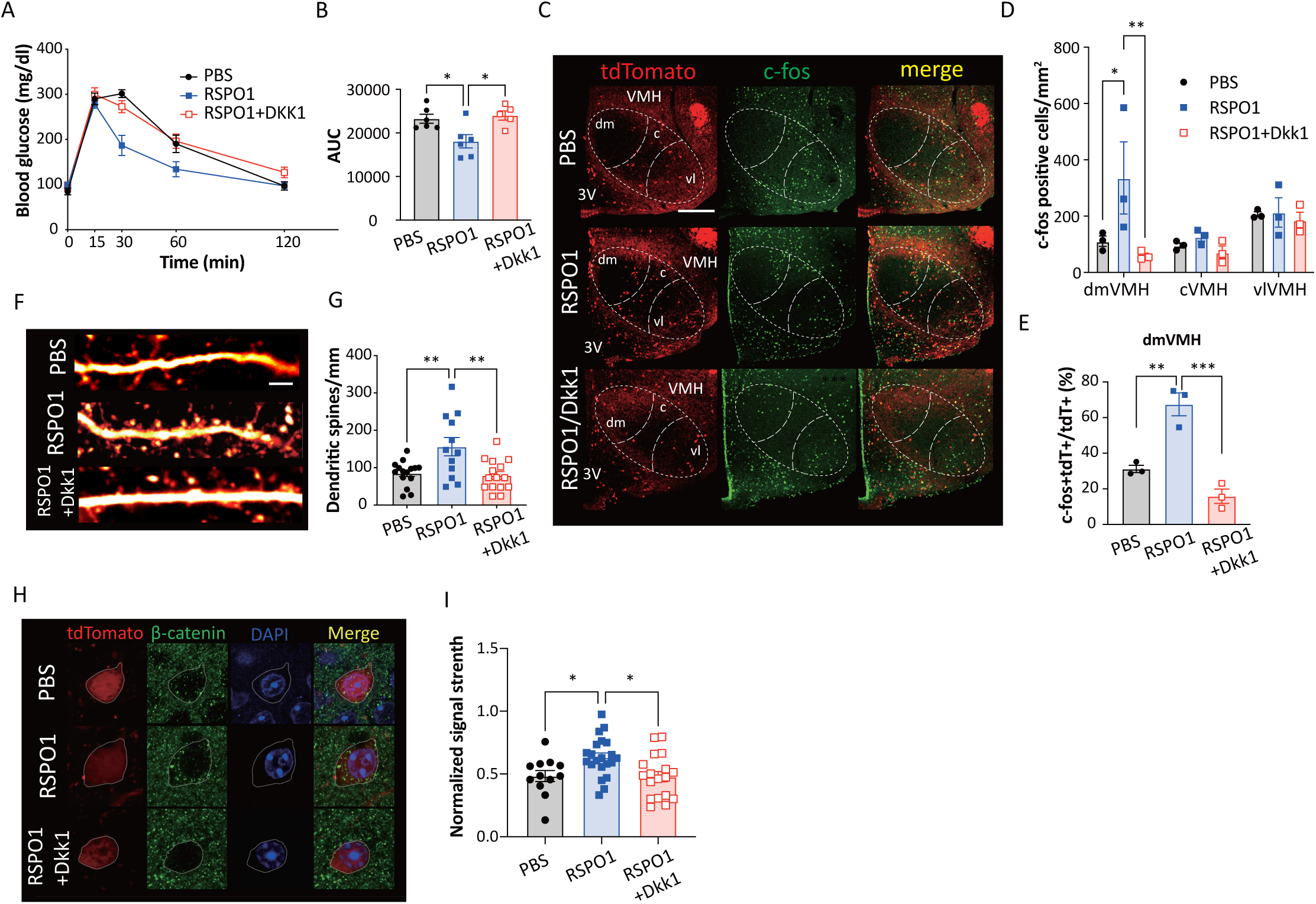
RSPO1 restores VMH^GE^ function in HFD-fed mice in a Wnt-dependent manner. (A and B) Effect of Wnt inhibition on RSPO1-mediated metabolic improvements. (A) Time course of blood glucose levels during GTT in HFD-fed mice treated with i.c.v. RSPO1 alone or in combination with the Wnt inhibitor Dkk1. (B) Area under the curve (AUC). (C–E) Assessment of glucose-induced c-fos expression in the VMH. (C) Representative micrographs of c-fos expression following glucose injection in each treatment group. Scale bar, 200μm. (D) Quantification of c-fos-positive cells in each VMH subregion. (E) Proportion of c-fos/tdTomato double-positive cells in the dmVMH. (F and G) Impact of Wnt modulation on dendritic spine density in VMH^GE^ neurons. (F) Representative micrographs of dendritic shafts and spines in VMH^GE^ neurons. Scale bar, 5μm. (G) Quantification of dendritic spine density (PBS: n = 15 cells from 3 mice; RSPO1: n = 12 cells from 3 mice; RSPO1+Dkk1: n = 15 cells from 3 mice). (H and I) Verification of Wnt signaling activation. (H) Representative immunohistochemistry images showing β-catenin expression in VMH^GE^ neurons following i.c.v. injection of RSPO1 or RSPO1+Dkk1. (I) Quantification of β-catenin signal intensity in VMH^GE^ neurons (PBS: n = 15 cells from 3 mice; RSPO1: n = 12 cells from 3 mice; RSPO1+Dkk1: n = 15 cells from 3 mice). All data represent the mean ± SEM; * *p*<0.05; ** *p*<0.01; *** *p*<0.001.

## Discussion

In this study, we labeled GE neurons *in vivo* using TRAP and confirmed their glucose responsiveness. By using this mouse model, we found that VMH^GE^ neurons are essential for regulating acute systemic glucose metabolism. This function is compromised by HFD feeding due to the inhibition of canonical Wnt signaling. Inhibition of Wnt signaling by HFD leads to synaptic loss (reduced dendritic spines) and blunted glucose-sensing in these neurons. We discovered that the injection of RSPO1, a Wnt enhancer downregulated by HFD, restores Wnt/β-catenin signaling and promotes spinogenesis. Consequently, central RSPO1 administration recovers neuronal glucose-sensing and improves systemic glucose tolerance by enhancing peripheral glucose utilization, identifying the RSPO1-Wnt axis as a key regulator of metabolic homeostasis.

The molecular machinery governing glucose sensing in VMH^GE^ neurons has long been considered analogous to that of pancreatic β-cells. Both cell types utilize glucose transporters and glucokinase to couple glycolysis with ATP-sensitive potassium (K_ATP_) channel closure and membrane depolarization^2,27^. Furthermore, mitochondrial function serves as a critical determinant of this metabolic coupling in both systems. In particular, Uncoupling Protein 2 (UCP2) plays a pivotal role in tuning the glucose-sensing of VMH neurons. UCP2 facilitates neuronal activation in the VMH and improves systemic glucose tolerance, highlighting the dependence of these neurons on robust mitochondrial bioenergetics for accurate glucose detection^9^. In line with this, our RNA-seq analysis revealed that HFD feeding markedly downregulated mitochondrial genes in VMH^GE^ neurons, which would impair glucose sensing of VMH^GE^ neurons.

Our previous study demonstrated that impairing neuronal glucose sensing in the dmVMH via cytosolic phospholipase A2 (cPLA2) knockdown significantly compromised glucose metabolism without affecting basal blood glucose levels or body weight^28^. Similarly, DREADD-mediated excitation of dmVMH SF1 neurons revealed that dmVMH activity does not alter basal blood glucose levels^6^. Furthermore, increasing the population of VMH^GE^ neurons through UCP2 overexpression improved glucose tolerance and insulin sensitivity while leaving body weight and energy expenditure unchanged^9^. These findings collectively suggest that VMH^GE^ neurons, particularly those in the dmVMH, do not regulate long-term energy homeostasis. Consistent with this, our specific manipulation of VMH^GE^ neurons altered glucose tolerance and insulin sensitivity without affecting basal blood glucose levels or body weight. Therefore, we conclude that VMH^GE^ neurons are specialized to control glucose utilization in response to acute fluctuations in systemic glucose levels, rather than maintaining long-term energy homeostasis. This functional distinction stands in contrast to POMC neurons in the arcuate nucleus, where mitochondrial function plays a pivotal role in long-term weight regulation. In POMC neurons, mitochondrial ROS production and UCP2-mediated uncoupling are essential for sensing systemic energy status to regulate body weight and satiety^29^. This is consistent with the finding that HFD-induced mitochondrial dysfunction disrupts the activity of glucose-sensing in POMC neurons^30^. Therefore, while mitochondrial impairment appears to be a common cellular pathology induced by HFD, its physiological consequence manifests differentially: driving obesity via POMC neurons, while causing “blindness” to acute glucose excursions in VMH^GE^ neurons.

Recently, emerging evidence indicates that β-cell populations are not functionally uniform but are orchestrated by specialized “leader” or “hub” cells—highly metabolically active and connected populations that dictate the collective response to glucose^31,32^. Analogous to these peripheral pacemakers, we propose that VMH^GE^ neurons function as the “hub nodes” of the central metabolic network. By integrating vast excitatory synaptic inputs via their dendritic spines, these neurons likely synchronize hypothalamic activity to efficiently transmit glucose information to peripheral organs. In this context, the HFD-induced loss of dendritic spines represents a selective functional disconnection of these hubs, thereby collapsing the coordinated output required for systemic glucose homeostasis.

Crucially, we identified canonical Wnt signaling as the key molecular guardian of this hub function, operating in a highly subregion-specific manner. Inhibition of Wnt signaling by Dkk1 suppressed glucose-induced c-fos expression exclusively in the dmVMH, while RSPO1 administration enhanced glucose sensing specifically in the same subregion. These results suggest that glucose sensing in the dmVMH is preferentially regulated by Wnt signaling. Functionally, the enhancement of dmVMH function by RSPO1 increased systemic insulin sensitivity by promoting glucose utilization rather than suppressing endogenous glucose production (EGP), which agrees with previous studies^28^. Mechanistically, beyond its established roles in neuronal structure and development, Wnt signaling also improves cellular glucose metabolism and ATP production by enhancing Akt activation, hexokinase activity, and the glycolytic rate^16,33^. Consistent with this, RSPO1 administration enhanced hypothalamic glucose uptake, indicating improved glucose metabolism in the hypothalamus of HFD-fed mice. Since ATP production is critical for VMH glucose sensing, increased hypothalamic glucose utilization likely represents a key step in recovering the glucose sensing capacity of VMH^GE^. Therefore, in contrast to the cVMH or vlVMH, Wnt signaling specifically regulates neuronal functions in the dmVMH, by improving cellular glucose metabolism. This further suggests that the VMH utilizes distinct mechanisms to detect ambient glucose levels across different subregions.

Transcriptomic analysis of VMH^GE^ revealed upregulated mRNA expression of excitatory postsynaptic genes. However, despite this increase in mRNA, dendritic spine density was reduced following HFD feeding. A likely explanation for this discrepancy is a blocking of mRNA translation. A similar translational block characterized by elevated mRNA transcription coupled with inhibited protein translation has been observed in β cell insulin production under HFD conditions^18,19^. Consistent with this, we found that HFD increased *Dlg4* mRNA in VMH^GE^ neurons while simultaneously decreasing hypothalamic PSD-95 protein levels. Similarly, β-catenin levels in VMH^GE^ were decreased despite elevated *Ctnnb1* mRNA expression following HFD feeding, further suggesting a translational blocking. The β-catenin is a core protein in canonical wnt signaling, which has been widely studied in regulation growth of axon, dendrite, and synaptogenesis^34,35^. Additionally, Wnt signaling also regulates ribosome biogenesis, which is impaired by HFD^35,36^ (SFig 4a), further supporting the translational block hypothesis.

Temporary downregulation of Wnt signaling reduces dendritic complexity and spine density, leading to abnormal behavior in adult rats^38^. Although the specific effects of Wnt signaling on hypothalamic synapses have been unclear, overexpression of GSK3β in the ARC impairs glucose metabolism, highlighting the importance of Wnt signaling in the hypothalamus^39^. Nevertheless, the specific role of Wnt signaling in the VMH remained undefined, despite the high expression of Wnt-related genes in this region^40^. Beyond synaptogenesis, Wnt signaling also regulates neuronal glucose metabolism by increasing ATP production, which directly contributes to glucose-sensing function^16,40,41^.

The reduction in dendritic spine density caused by Dkk1 implies that Wnt signaling is essential for maintaining the structural and functional integrity of VMH^GE^ neurons. Similarly, HFD feeding also resulted in a loss of dendritic spines in this population. Synaptic stimulation has been shown to transiently boost glycolysis and increase cytosolic NADH levels^42^. This increased glycolytic flux likely enhances intracellular ATP production, thereby facilitating glucose sensing. This suggests a reciprocal relationship between cellular glucose metabolism and neuronal activity, indicating that synaptic stimulation regulates glucose sensing via increased intracellular energy production. Consistent with this, the blockade of AMPA and NMDA receptors blunted glucose-induced activity in VMH^GE^. Indeed, VMH^GE^ glucose sensing is known to rely, at least in part, on afferent projections^43^. In summary, VMH^GE^ regulate acute glucose metabolism in response to abrupt fluctuations in blood glucose, but do not appear to control long-term energy homeostasis. This critical function is maintained by Wnt signaling, which is inhibited by HFD feeding.

In summary, our study unveils a critical link between diet-induced synaptic remodeling and systemic metabolic dysfunction. We demonstrate that the impairment of VMH^GE^ glucose sensing under obesity is a structural failure driven by the suppression of the RSPO1-Wnt signaling axis. Importantly, restoring this pathway is sufficient to rebuild dendritic spines and recover metabolic control. Consequently, our findings propose a paradigm shift in understanding metabolic diseases: targeting the structural plasticity of hypothalamic circuits—specifically via Wnt enhancers like RSPO1—offers a novel and promising therapeutic strategy to combat the glucose intolerance associated with type 2 diabetes and obesity.

## Author contributions

M.L. and C.T. designed the research project. M.L., C.C., S.H., and T.A. performed the experiments. M.L. and C.T. performed the data analysis. R.E. and C.T. supervised the research. All authors contributed to the writing of the manuscript and approved the submitted version.

## Supporting information

graphical abstract

---

**Supplementary Figure 1.**
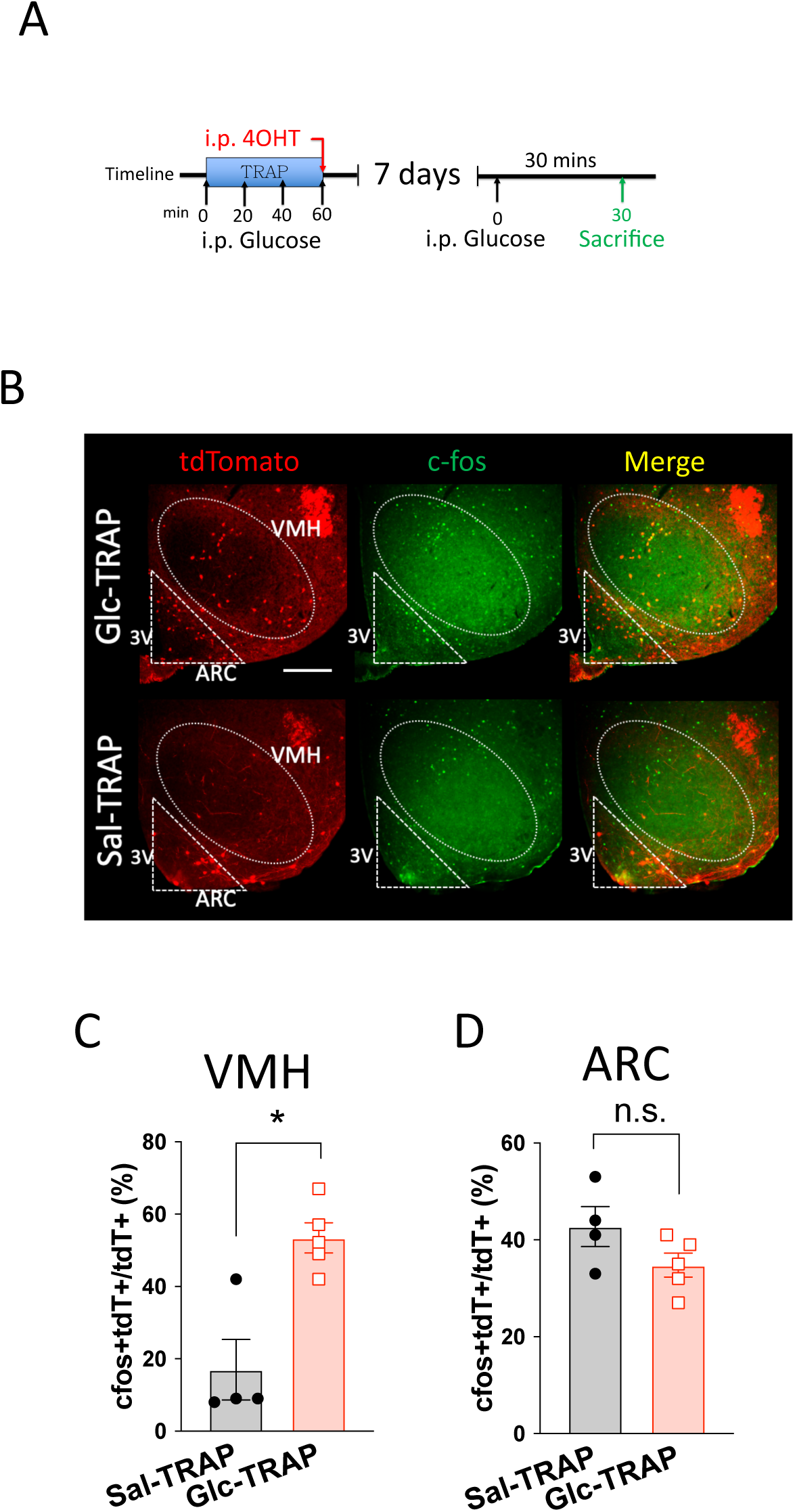
Glucose responsiveness of Glc-TRAPed neurons in the VMH and ARC. (A) Schematic of the experimental design for TRAP-labeling GE neurons and assessing their reactivation by glucose using c-fos expression. (B–D) Assessment of glucose-induced c-fos expression in Glc-TRAP and Sal-TRAP mice. (B) Representative micrographs showing c-fos immunoreactivity and TRAP-labeled (tdTomato+) neurons following glucose injection. Scale bar, 200 μm. (C and D) Proportion of c-fos/tdTomato double-positive cells among the TRAP-labeled population in the VMH (C) and ARC (D). All data represent the mean ± SEM; * *p*<0.05.

**Supplementary Figure 2.**
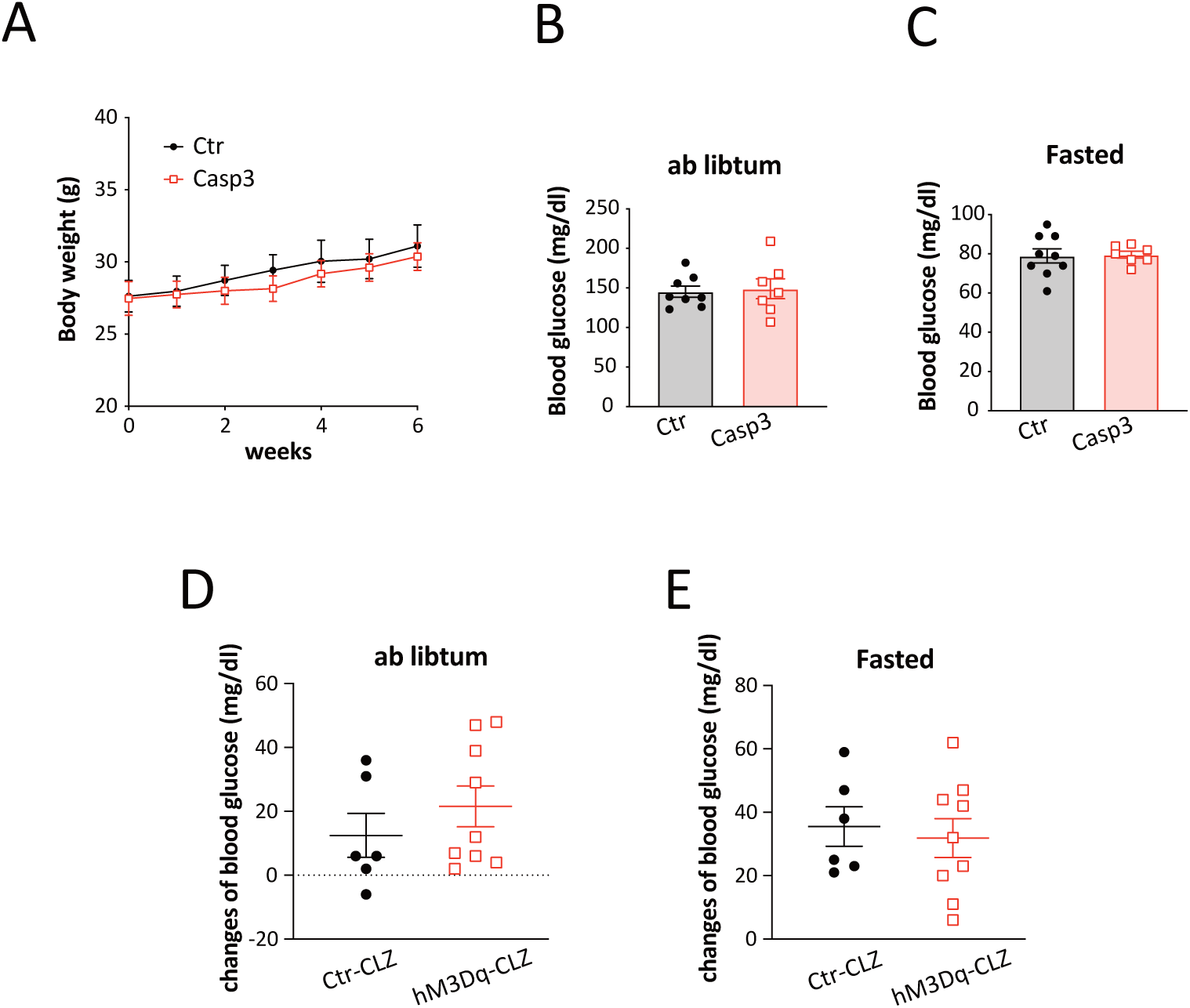
Effects of Ablation or chemogenetic activation of VMH^GE^ on long-term glucose homeostasis. (A) Body weight of control and VMH^GE^-ablated mice measured for six weeks after 4OHT induction. (Control: n = 4, Caspase-3: n = 4). (B and C) Basal blood glucose levels 8 weeks after TRAP induction. (B) *Ad libitum* fed state and (C) fasted state. (D and E) Acute effect of chemogenetic activation of VMH^GE^ neurons on base-line blood glucose levels. Blood glucose was measured 30 min following clozapine injection under (D) *ad libitum* fed or (E) fasted conditions. All data represent the mean ± SEM.

**Supplementary Figure 3.**
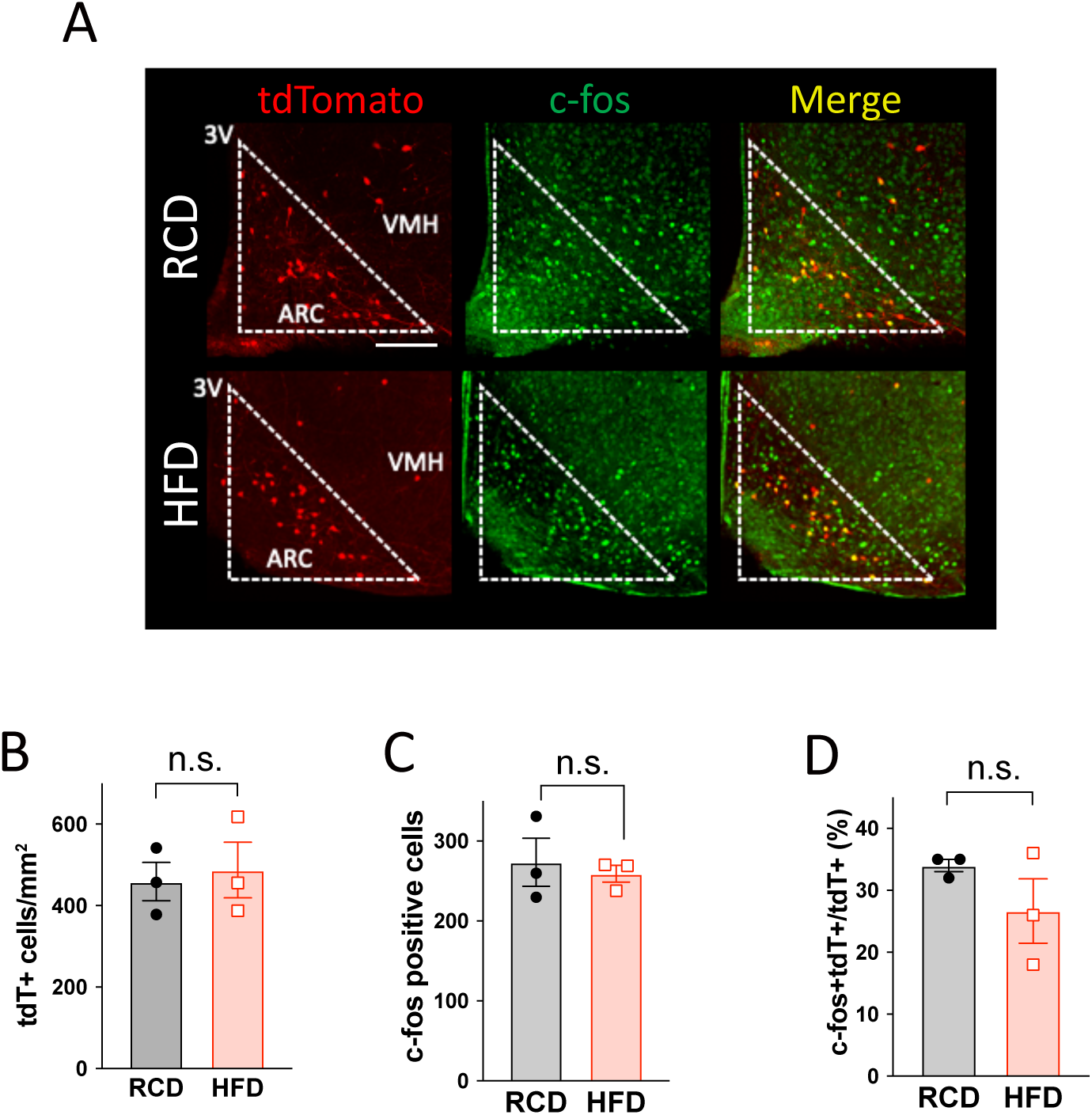
HFD feeding does not affect the population of glucose responsiveness in the ARC. (A) Representative micrographs of c-fos immunostaining in the arcuate nucleus (ARC) of Glc-TRAP mice fed RCD or HFD. Scale bar, 100 μm. (B–D) Quantification of neuronal populations in the ARC. (B) Number of TRAP-labeled (tdT+) neurons. (C) Number of c-fos-positive cells. (D) Proportion of c-fos/tdTomato double-positive cells. All data represent the mean ± SEM.

**Supplementary Figure 4.**
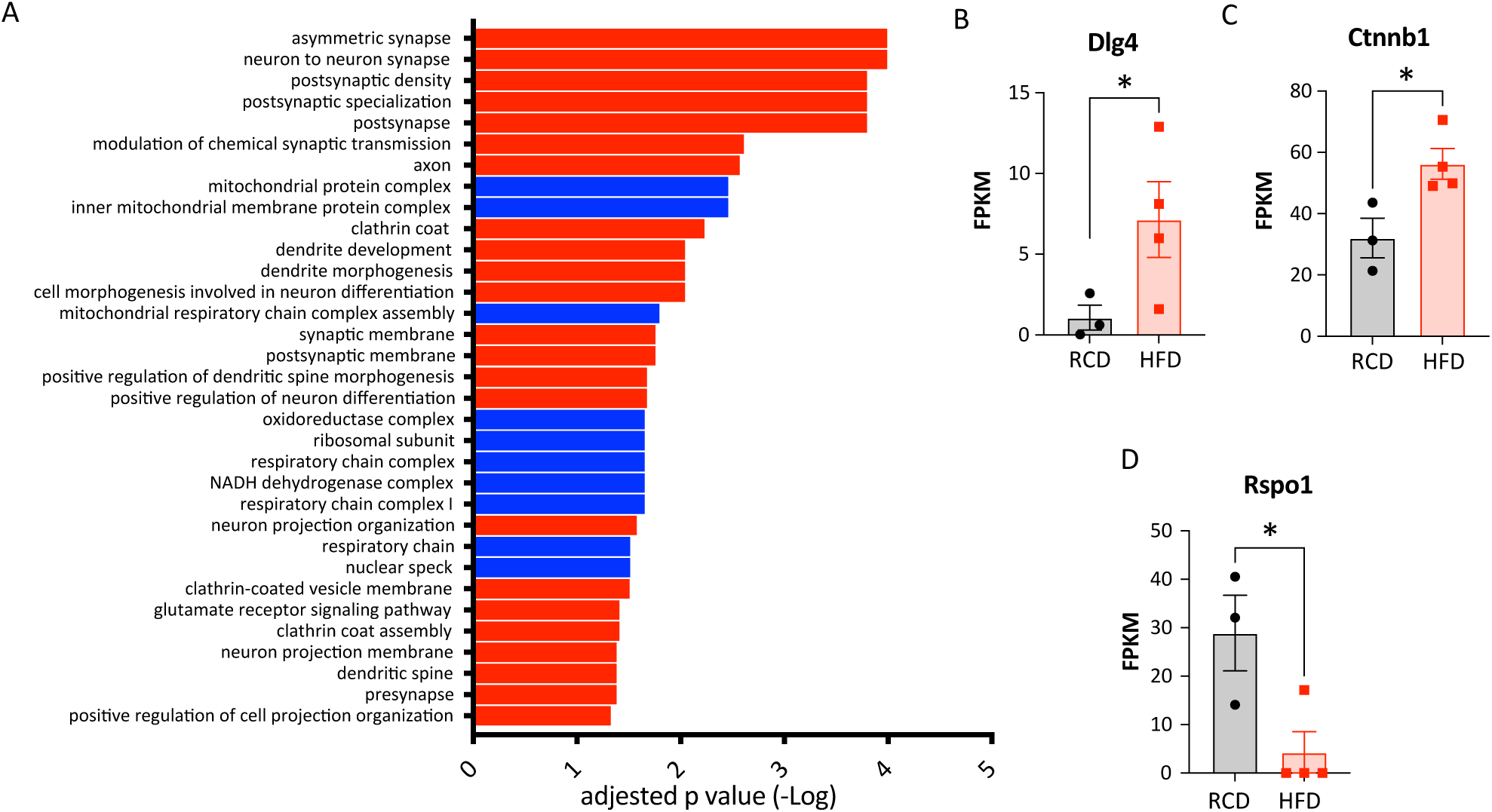
RNA sequencing of VMH^GE^ neurons from RCD and HFD fed mice. (A) Gene ontology enrichment analysis of differentially expressed genes (DEGs) in VMH^GE^ neurons from RCD-fed versus HFD-fed mice. Red bars represent upregulated gene sets and blue bars represent downregulated gene sets in VMH^GE^ neurons after 3 months of HFD feeding. (B–D) mRNA expression levels of (B) *Dlg4* (PSD95), (C) *Ctnnb1* (β-catenin), and (D) *Rspo1* derived from the RNA-seq dataset. All data represent the mean ± SEM; * *p*<0.05.

**Supplementary Figure 5.**
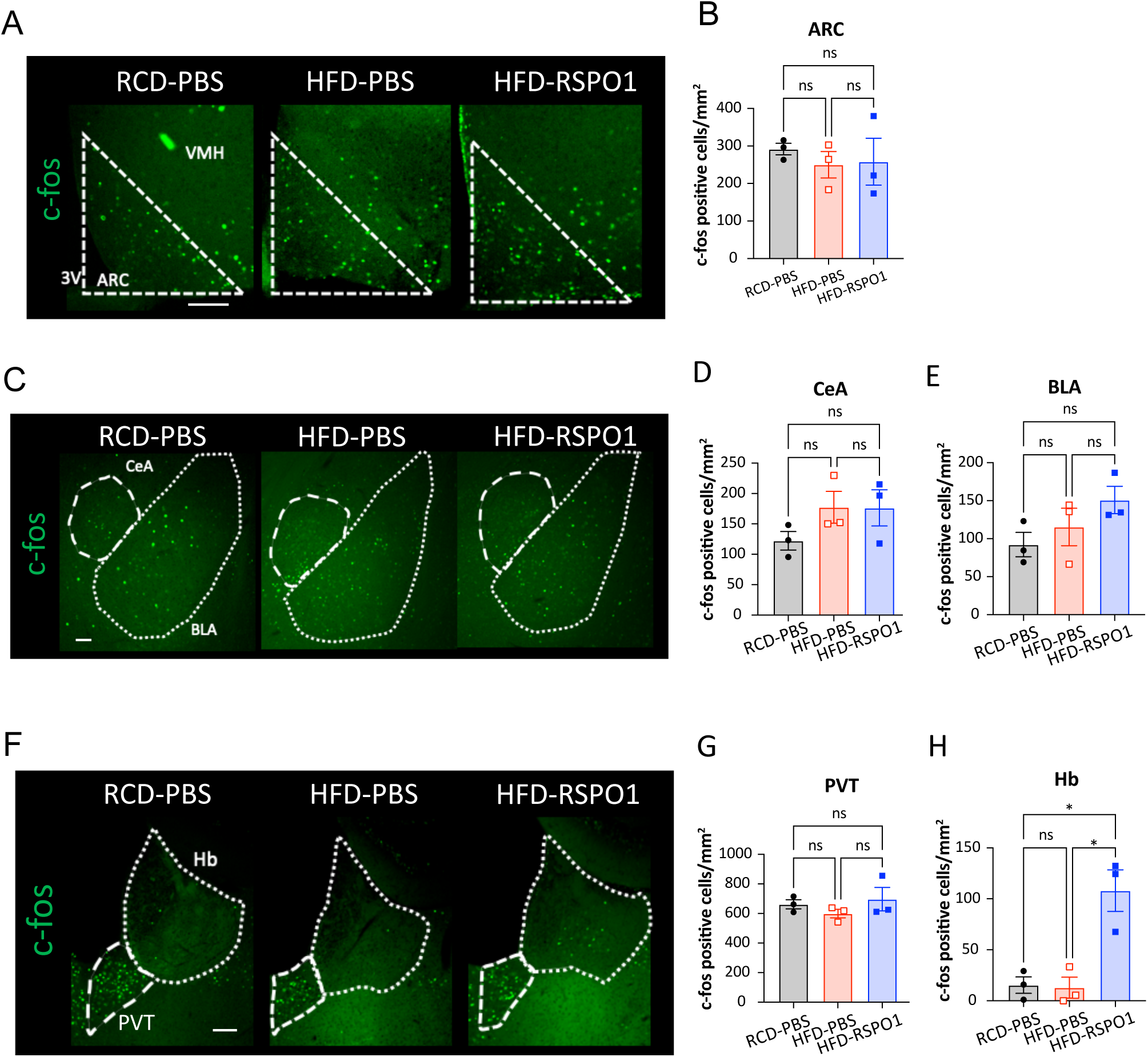
Effects of RSPO1 on glucose-induced c-fos expression in LGR4-expressing regions in RCD- or HFD-fed mice. (A and B) Effect of RSPO1 on the arcuate nucleus (ARC). (A) Representative micrographs of c-fos immunostaining in the ARC following i.c.v RSPO1 and i.p. glucose injection. Scale bar, 100 μm. (B) Number of c-fos-positive cells in the ARC. (C–H) Effect of RSPO1 on the central (CeA), basolateral amygdala (BLA), habenula (Hb) and paraventricular thalamus (PVT). (C) Representative micrographs of c-fos immunostaining in the amygdala. Scale bar, 100 μm. (D-E) Number of c-fos-positive cells in the CeA (D) and BLA (E) following i.c.v RSPO1 and i.p. glucose injection. (F) Number of c-fos-positive cells in the Hb and PVT. Scale bar, 100 μm. (G-H) Number of c-fos-positive cells in the Hb (G) and PVT (H) . All data represent the mean ± SEM;** *p*<0.01.

**Supplementary Figure 6.**
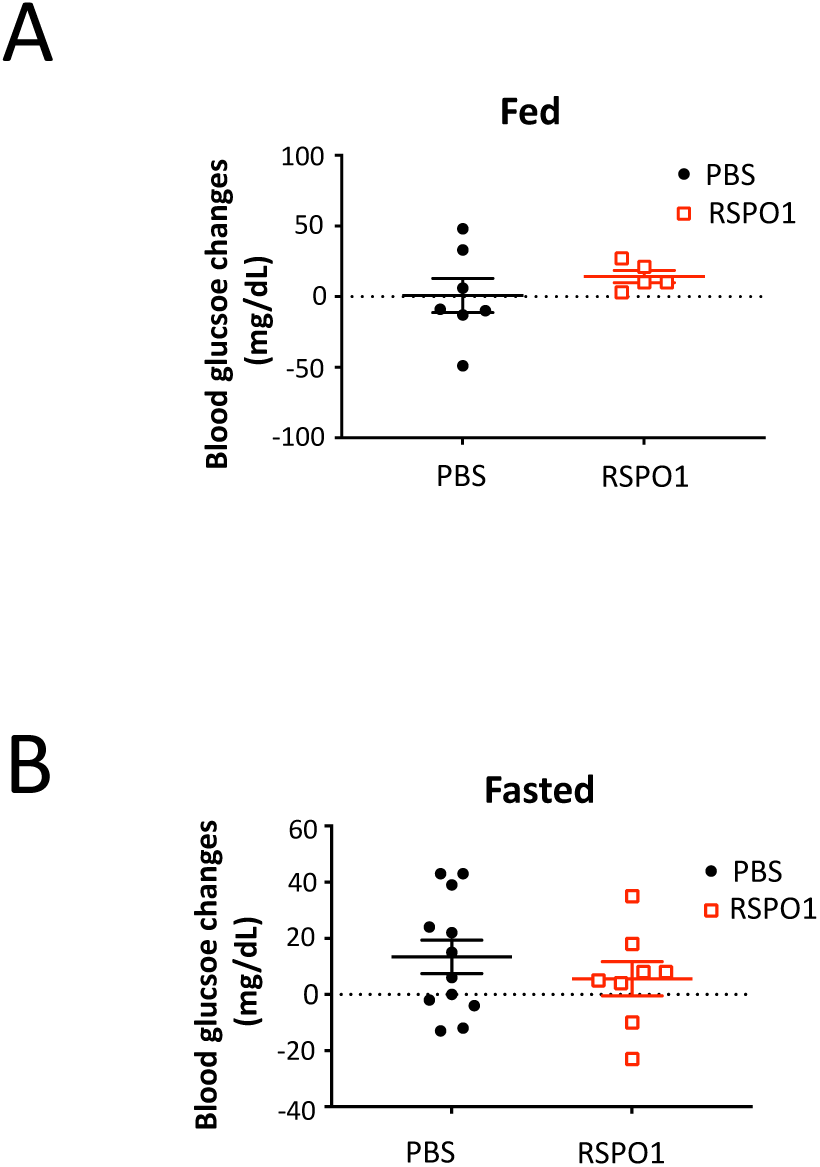
Impact of RSPO1 administration on basal blood glucose levels. (A and B) Basal blood glucose levels in HFD-fed mice 3 h following i.c.v. injection of RSPO1. (A) *Ad libitum* fed state. (B) Fasted state. All data represent the mean ± SEM.

## Reference

1. Kleinridders, A. et al. Regional differences in brain glucose metabolism determined by imaging mass spectrometry. Mol. Metab. 12, 113–121 (2018).

2. Routh, V. H. Glucose sensing neurons in the ventromedial hypothalamus. Sensors 10, 9002–9025 (2010).

3. Shimazu, T. & Minokoshi, Y. Systemic Glucoregulation by Glucose-Sensing Neurons in the Ventromedial Hypothalamic Nucleus (VMH). J. Endocr. Soc. 1, 449–459 (2017).

4. Meek, T. H., et al. Functional identification of a neurocircuit regulating blood glucose. Proc. Natl. Acad. Sci. 113, E2073–E2082 (2016).

5. He, Y. et al. Estrogen receptor-α expressing neurons in the ventrolateral VMH regulate glucose balance. Nat. Commun. 11, 2165 (2020).

6. Coutinho, E. A. et al. Activation of SF1 Neurons in the Ventromedial Hypothalamus by DREADD Technology Increases Insulin Sensitivity in Peripheral Tissues. Diabetes 66, 2372–2386 (2017).

7. Burdakov, D., Luckman, S. M. & Verkhratsky, A. Glucose-sensing neurons of the hypothalamus. Philos. Trans. R. Soc. Lond. B. Biol. Sci. 360, 2227–2235 (2005).

8. Routh, V. H., Hao, L., Santiago, A. M., Sheng, Z. & Zhou, C. Hypothalamic glucose sensing: making ends meet. Front. Syst. Neurosci. 8, (2014).

9. Toda, C. et al. UCP2 Regulates Mitochondrial Fission and Ventromedial Nucleus Control of Glucose Responsiveness. Cell 164, 872–883 (2016).

10. Sethi, J. K. & Vidal-Puig, A. Wnt signalling and the control of cellular metabolism. Biochem. J. 427, 1–17 (2010).

11. Grant, S. F. A. et al. Variant of transcription factor 7-like 2 (TCF7L2) gene confers risk of type 2 diabetes. Nat. Genet. 38, 320–323 (2006).

12. Ip, W., Chiang, Y.-T. A. & Jin, T. The involvement of the wnt signaling pathway and TCF7L2 in diabetes mellitus: The current understanding, dispute, and perspective. Cell Biosci. 2, 28 (2012).

13. He, C.-W., Liao, C.-P. & Pan, C.-L. Wnt signalling in the development of axon, dendrites and synapses. Open Biol. 8, 180116 (2018).

14. Hussaini, S. M. Q. et al. Wnt signaling in neuropsychiatric disorders: Ties with adult hippocampal neurogenesis and behavior. Neurosci. Biobehav. Rev. 47, 369–383 (2014).

15. Zou, Y. Wnt signaling in axon guidance. Trends Neurosci. 27, 528–532 (2004).

16. Villaseca, P., Cisternas, P. & Inestrosa, N. C. Menopause and development of Alzheimer’s disease: Roles of neural glucose metabolism and Wnt signaling. Front. Endocrinol. 13, (2022).

17. Guenthner, C. J., Miyamichi, K., Yang, H. H., Heller, H. C. & Luo, L. Permanent genetic access to transiently active neurons via TRAP: targeted recombination in active populations. Neuron 78, 773–784 (2013).

18. Hatanaka, M. et al. Chronic high fat feeding restricts islet mRNA translation initiation independently of ER stress via DNA damage and p53 activation. Sci. Rep. 7, 3758 (2017).

19. Li, Z. et al. RNA-binding protein DDX1 is responsible for fatty acid-mediated repression of insulin translation. Nucleic Acids Res. 46, 12052–12066 (2018).

20. Ebrahimi, K. B. et al. Oxidative Stress Induces an Interactive Decline in Wnt and Nrf2 Signaling in Degenerating Retinal Pigment Epithelium. Antioxid. Redox Signal. 29, 389–407 (2018).

21. Palomera-Avalos, V. et al. Resveratrol Protects SAMP8 Brain Under Metabolic Stress: Focus on Mitochondrial Function and Wnt Pathway. Mol. Neurobiol. 54, 1661–1676 (2017).

22. Purro, S. A., Dickins, E. M. & Salinas, P. C. The secreted Wnt antagonist Dickkopf-1 is required for amyloid β-mediated synaptic loss. J. Neurosci. Off. J. Soc. Neurosci. 32, 3492–3498 (2012).

23. Li, J.-Y. et al. LGR4 and its ligands, R-spondin 1 and R-spondin 3, regulate food intake in the hypothalamus of male rats. Endocrinology 155, 429–440 (2014).

24. Glinka, A. et al. LGR4 and LGR5 are R-spondin receptors mediating Wnt/β-catenin and Wnt/PCP signalling. EMBO Rep. 12, 1055–1061 (2011).

25. Hao, H.-X., Jiang, X. & Cong, F. Control of Wnt Receptor Turnover by R-spondin-ZNRF3/RNF43 Signaling Module and Its Dysregulation in Cancer. Cancers 8, 54 (2016).

26. Li, J.-Y., Wu, X., Lee, A., Zhou, S.-Y. & Owyang, C. Altered R-spondin 1/CART neurocircuit in the hypothalamus contributes to hyperphagia in diabetes. J. Neurophysiol. 121, 928–939 (2019).

27. Jin, E., Briggs, J. K., Benninger, R. K. & Merrins, M. J. Glucokinase activity controls peripherally-located subpopulations of β-cells that lead islet Ca2+ oscillations. eLife 13, (2025).

28. Lee, M.-L. et al. Prostaglandin in the ventromedial hypothalamus regulates peripheral glucose metabolism. Nat. Commun. 12, 2330 (2021).

29. Andrews, Z. B. et al. UCP2 mediates ghrelin’s action on NPY/AgRP neurons by lowering free radicals. Nature 454, 846–851 (2008).

30. Parton, L. E. et al. Glucose sensing by POMC neurons regulates glucose homeostasis and is impaired in obesity. Nature 449, 228–232 (2007).

31. Slak Rupnik, M. Opportunity makes a hub or a leader. eLife 14, e105929 (2025).

32. Johnston, N. R. et al. Beta Cell Hubs Dictate Pancreatic Islet Responses to Glucose. Cell Metab. 24, 389–401 (2016).

33. Cisternas, P., Salazar, P., Silva-Álvarez, C., Barros, L. F. & Inestrosa, N. C. Activation of Wnt Signaling in Cortical Neurons Enhances Glucose Utilization through Glycolysis. J. Biol. Chem. 291, 25950–25964 (2016).

34. McLeod, F. & Salinas, P. C. Wnt proteins as modulators of synaptic plasticity. Curr. Opin. Neurobiol. 53, 90–95 (2018).

35. Rosso, S. B. & Inestrosa, N. C. WNT signaling in neuronal maturation and synaptogenesis. Front. Cell. Neurosci. 7, 103 (2013).

36. Madan, B. et al. Temporal dynamics of Wnt-dependent transcriptome reveal an oncogenic Wnt/MYC/ribosome axis. J. Clin. Invest. 128, 5620–5633 (2019).

37. Pfister, A. S. & Kühl, M. Of Wnts and Ribosomes. Prog. Mol. Biol. Transl. Sci. 153, 131–155 (2018).

38. Viale, B. et al. Transient Deregulation of Canonical Wnt Signaling in Developing Pyramidal Neurons Leads to Dendritic Defects and Impaired Behavior. Cell Rep. 27, 1487–1502.e6 (2019).

39. Benzler, J. et al. Hypothalamic glycogen synthase kinase 3β has a central role in the regulation of food intake and glucose metabolism. Biochem. J. 447, 175–184 (2012).

40. Nusse, R. & Clevers, H. Wnt/β-Catenin Signaling, Disease, and Emerging Therapeutic Modalities. Cell 169, 985–999 (2017).

41. Cisternas, P. et al. Wnt-induced activation of glucose metabolism mediates the in vivo neuroprotective roles of Wnt signaling in Alzheimer disease. J. Neurochem. 149, 54–72 (2019).

42. Díaz-García, C. M. et al. Neuronal Stimulation Triggers Neuronal Glycolysis and Not Lactate Uptake. Cell Metab. 26, 361–374.e4 (2017).

43. Song, Z., Levin, B. E., McArdle, J. J., Bakhos, N. & Routh, V. H. Convergence of Pre- and Postsynaptic Influences on Glucosensing Neurons in the Ventromedial Hypothalamic Nucleus. Diabetes 50, 2673–2681 (2001).

