## Supplementary figures and images for "R-spondin 1 restores hypothalamic glucose-sensing and systemic glucose homeostasis via Wnt signaling in diet-induced obese mice"

### graphical abstract

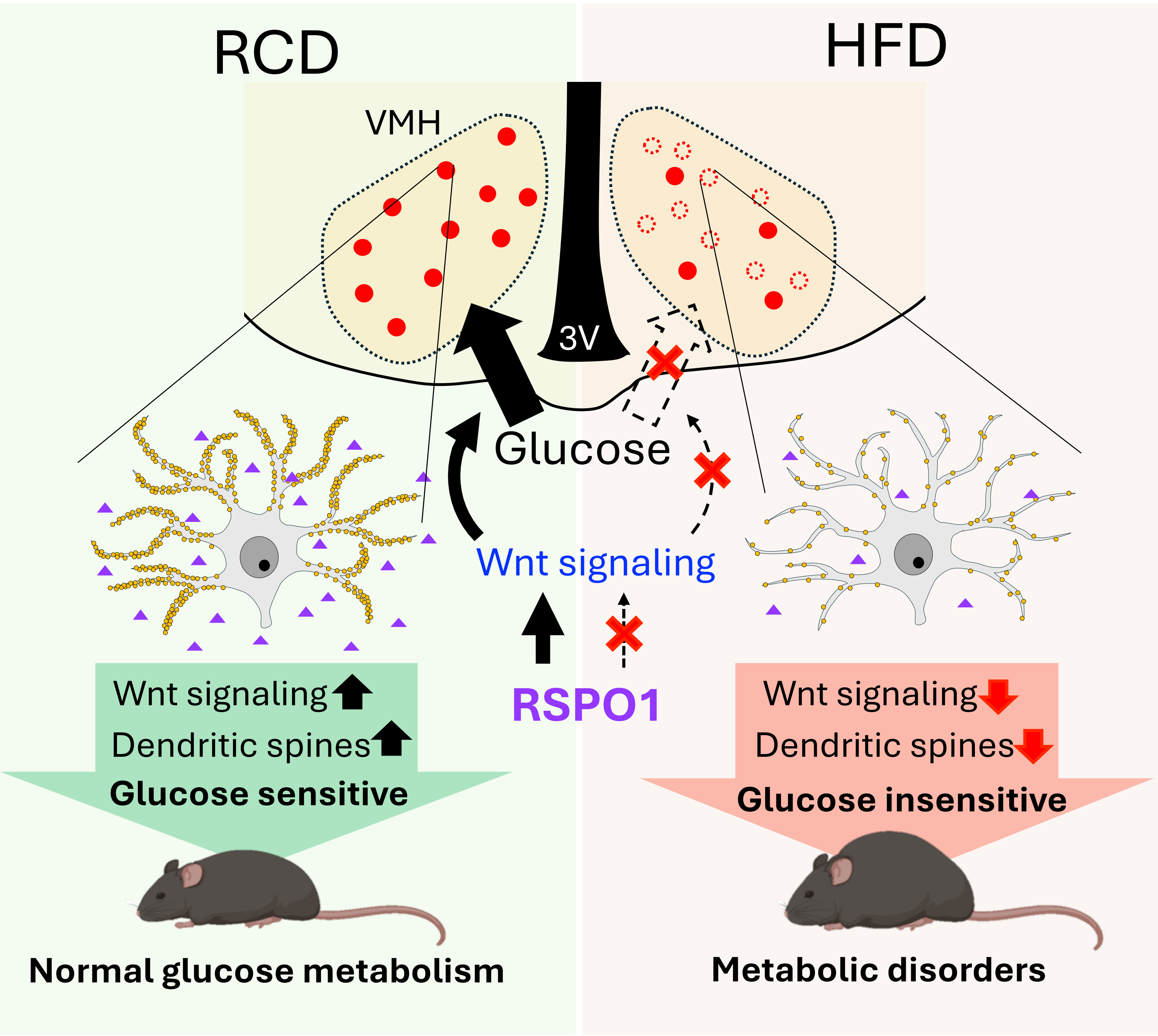
