## Supplementary material for "R-spondin 1 restores hypothalamic glucose-sensing and systemic glucose homeostasis via Wnt signaling in diet-induced obese mice": method

### STAR Methods

#### RESOURCE AVAILABILITY

##### Lead contact

##### Materials availability

This study did not generate new unique reagents

#### Data and code availability

All data needed to evaluate the conclusions in the study are presented in the main text and Supplementary Materials. Further study-related data can be requested from the authors.

#### EXPERIMENTAL MODEL AND SUBJECT DETAILS

##### Mouse model

Arc-cre$^{ER}$ and Ai14 mice were purchased from the Jackson laboratory (Strain #021881 and 007914; Bar Harbor, ME). These two strains were crossed to generate [Arc-cre$^{ER}$]::Ai14 double-transgenic mice. For hyperinsulinemic-euglycemic clamp studies, male C57BL/6J mice were purchased from Charles River Laboratories Japan. All mice were housed under controlled temperature (22–24°C) and a 12-h light/12-h dark cycle, with *ad libitum* access to food and water. All animal care and experimental procedures were performed in accordance with the guidelines and approval of the Animal Care and Use Committees of Hokkaido University, Kumamoto University, and the National Institute for Physiological Sciences.

#### METHOD DETAILS

##### TRAP experiments

4-Hydroxytamoxifen (4-OHT; H6278, Sigma-Aldrich) was dissolved in DMSO at a concentration of 100 mg/mL and stored at −80°C until use. On the day of the TRAP experiments, the 100 mg/mL 4-OHT stock solution was diluted 5-fold with DMSO, followed by a 20-fold dilution in saline containing 3% Tween 80. The final diluted 4-OHT solution (1 mg/mL) was kept at 4°C for no longer than 3 h. Eight- to twelve-week-old male mice received four injections of glucose and a single injection of 4-OHT via intraperitoneal (i.p.) injection over a 60-min period. Specifically, glucose (3 g/kg in saline) was administered at 20-min intervals (t = 0, 20, 40, and 60 min), and 4-OHT (10 mg/kg) was injected immediately after the fourth glucose injection (t = 60 min). To trigger the expression of AAV-delivered genes (e.g., Caspase-3, hM3Dq, and GCaMP6s), TRAP induction was performed 1 week after viral injection. All TRAP experiments were conducted between 14:00 and 15:00 to minimize the confounding effects of circadian rhythms.

**Immunohistchemistry**

Mice were transcardially perfused with heparinized saline followed by 4% paraformaldehyde (PFA). To assess the glucose responsiveness of VMH neurons via cfos expression, animals received an i.p. injection of glucose (3 g/kg) 30 min prior to perfusion. For evaluating the effects of RSPO1 and/or Dkk1 on neuronal activity, recombinant RSPO1 protein (1 μg in 0.5 μL; Cat# 3474-RS, R&D Systems, MN) and/or Dkk1 (100 ng in 0.5 μL; Cat# 5897-DK, R&D Systems, MN) were intracerebroventricularly (i.c.v.) injected 3 h before glucose administration. Brains were post-fixed in 4% PFA overnight, and 50-μm-thick coronal sections containing the entire VMH were prepared using a vibratome. Free-floating sections were incubated with a rabbit anti-cfos antibody (1:2000; Cell Signaling Technology, MA) or a rabbit anti-β-catenin antibody (1:250; sc-7963, Santa Cruz) in a staining solution (0.1 M phosphate buffer [PB] containing 4% normal goat serum, 0.1% glycine, and 0.2% Triton X-100) overnight at room temperature. After washing with 0.1 M PB, the sections were incubated with an Alexa Fluor 488-conjugated goat anti-rabbit IgG secondary antibody (1:500; Cell Signaling Technology, MA) for 2 h at room temperature. The stained sections were rinsed several times with 0.1 M PB and mounted on glass slides using VECTASHIELD Antifade Mounting Medium (Vector Laboratories, CA).

**Stereotaxic surgeries and AAV injection**

Eight- to ten-week-old male mice were anesthetized with a mixture of ketamine (100 mg/kg) and xylazine (10 mg/kg) and placed in a stereotaxic frame (Narishige, Tokyo, Japan). For i.c.v. cannulation, guide cannulae were implanted into the lateral ventricle using the following coordinates relative to bregma: anterior-posterior (AP), −0.3 mm; mediolateral (ML), 1.0 mm; and dorsoventral (DV), −2.5 mm from the skull surface. The cannulae were secured to the skull using cyanoacrylate adhesive and dental cement. Mice were allowed to recover for 7 days prior to subsequent experiments.

For the ablation of $VMH^{GE}$ neurons, 0.3 μL of AAV5-flex-taCasp3-TEVp (1.0 × $10^{12}$ GC/ml; University of North Carolina Vector Core, NC) was bilaterally injected into the VMH of Arc-$Cre^{ER}$::Ai14 double-transgenic mice using the following coordinates: AP, −1.4 mm; ML, ±0.4 mm; and DV, −5.7 mm. For chemogenetic manipulation, 0.3 μL of AAV2-hSyn-DIO-hM3Dq-mCherry (1.0 × $10^{12}$ GC/ml; Cat#50474, Addgene, MA) or AAV2-hSyn-DIO-mCherry (1.0 × $10^{12}$ GC/ml; University of North Carolina Vector Core, NC) was injected into the VMH of Arc-Cre$^{ER}$ mice. For slice calcium imaging, AAV1-EF1a-DIO-GCaMP6s (1.0 × $10^{12}$ GC/ml; Cat#51082, Addgene, MA) or a 1:1 mixture of AAV1-EF1a-DIO-GCaMP6s and AAV2-hSyn-mCherry-Cre (1.0 × $10^{12}$ GC/ml; University of North Carolina Vector Core, NC) was injected into the VMH using the same coordinates. Following the injections, the skin incisions were closed with sutures.

**Glucose- and insulin-tolerance test**

Glucose tolerance tests (GTTs) were performed on either *ad libitum*-fed or 16-h fasted mice. The *ad libitum*-fed animals were used to assess the effects of Dkk1, whereas the fasted mice were used to evaluate VMH$^{GE}$ function and the effects of RSPO1. Mice received an intraperitoneal (i.p.) injection of glucose (2 g/kg body weight), and blood glucose levels were measured using a handheld glucometer (FreeStyle, Nipro, Japan) at 0, 15, 30, 60, and 120 min post-injection.

Insulin tolerance tests (ITTs) were conducted on *ad libitum*-fed mice. Animals received an i.p. injection of insulin (0.5 U/kg body weight; Novo Nordisk, Denmark) diluted in saline containing 0.2% bovine serum albumin (BSA). Blood glucose levels were monitored at 0, 15, 30, 60, and 120 min post-injection.

For chemogenetic (DREADD) experiments, clozapine (0.1 mg/kg body weight; Cayman Chemical, MI) was administered i.p. 30 min prior to the glucose or insulin challenge ($t = -30$ min). To evaluate the involvement of Wnt signaling, recombinant RSPO1 (1 μg in 0.5 μL; Cat# 3474-RS, R&D Systems, MN) and/or Dkk1 (100 ng in 0.5 μL; Cat# 5897-DK, R&D Systems, MN) were administered i.c.v. either alone or in combination 3 h prior to the GTT or ITT.

**Ex vivo calcium imaging**

One week following viral injection, mice injected with AAV1-EF1α-DIO-GCaMP6s underwent TRAP induction to trigger GCaMP6s expression in VMH$^{GE}$ neurons. Mice injected with the viral mixture (AAV1-EF1α-DIO-GCaMP6s and AAV2-hSyn-mCherry-Cre) did not undergo TRAP induction. Both groups were maintained on a regular chow diet (RCD) for an additional 12 weeks prior to the calcium imaging experiments. To assess the impact of an HFD on VMH$^{GE}$ function, a subset of TRAP-induced mice was randomly assigned to receive either an RCD or HFD for 12 weeks. To evaluate the effect of RSPO1 on glucose sensing, guide cannulae were implanted into the lateral ventricles of HFD-fed mice 1 week prior to imaging. On the day of the experiment, RSPO1 or PBS was administered i.c.v. 3 h prior to brain extraction.

Acute coronal brain slices from adult mice were prepared based on a previously described protocol[1] with minor modifications. Briefly, mice were transcardially perfused with ~15 mL of ice-cold NMDG-based artificial cerebrospinal fluid (aCSF) containing 92mM NMDG, 2.5mM KCl, 1.25mM $NaH_2PO_4$, 30mM $NaHCO_3$, 20mM HEPES, 2mM thiourea, 5mM sodium ascorbate, 3mM sodium pyruvate, 10mM $MgSO_4$, 0.5mM $CaCl_2$, 25mM glucose. Brain slices (250-μm thickness) were cut in ice-cold NMDG aCSF using a vibratome and incubated at 34°C for 13 min, followed by a gradual sodium spiking procedure as previously described[1]. The slices were then transferred to holding aCSF (92mM NaCl, 2.5mM KCl, 1.25mM $NaH_2PO_4$, 20mM HEPES, 2mM thiourea, 5mM sodium ascorbate, 3mM sodium pyruvate, 20mM $NaHCO_3$, 2mM $MgSO_4$, 2mM $CaCl_2$, 12.5mM glucose) and allowed to recover for 1 h at room temperature prior to recording. During imaging, individual slices were transferred to a recording chamber and continuously perfused with recording aCSF at a flow rate of 3 mL/min. To classify glucose-sensing neurons, brain slices were sequentially perfused with recording aCSF containing alternating glucose concentrations (2.5 mM, 0.2 mM, and back to 2.5 mM). Neurons exhibiting a >30% change in calcium fluorescence intensity in response to these glucose fluctuations were categorized as either glucose-excited or glucose-inhibited neurons. At the end of each recording session, KCl was bath-applied to induce neuronal depolarization, thereby confirming cell viability and functional calcium responsiveness.

**$VMH^{GE}$ Cell Sorting and RNA Sequencing**

Eight-week-old male Arc-$Cre^{ER}$::Ai14 double-transgenic mice underwent TRAP induction to label $VMH^{GE}$ neurons. The mice were then randomly assigned to receive either an RCD or an HFD for the subsequent 12 weeks. On the day of cell sorting, mice were anesthetized with isoflurane and decapitated. Brains were rapidly extracted and immediately submerged in ice-cold sucrose-based cutting solution (250 mM sucrose, 2.5 mM KCl, 6 mM $MgCl_2$, 1 mM $CaCl_2$, 1.25 mM $NaH_2PO_4$, 26 mM $NaHCO_3$, and 10 mM glucose; pH 7.4). Coronal brain slices (500-μm thickness) containing the entire VMH were prepared using a vibratome, and the VMH was microdissected using a clean surgical blade under a dissecting microscope. The VMH tissue explants were transferred to a trituration solution consisting of artificial cerebrospinal fluid (aCSF; 124 mM NaCl, 3 mM KCl, 2 mM $CaCl_2$, 2 mM $MgCl_2$, 1.23 mM $NaH_2PO_4$, 26 mM $NaHCO_3$, and 10 mM glucose) supplemented with 20 U/mL papain, 0.015 mg/mL DNase I, and 0.075 mg/mL BSA. Following a 15-min incubation at 34°C, the tissue was washed with aCSF and mechanically triturated using a fire-polished glass pipette, followed sequentially by 1-mL and 200-μL plastic pipette tips. The suspension was centrifuged at 100 × g for 5 min to remove large tissue debris. The resulting cell suspension in aCSF was plated onto a Petri dish and left undisturbed at room temperature for 1 h. tdTomato-positive cells were manually sorted using a glass capillary and transferred into clean PBS containing 0.075 mg/mL BSA. Batches of five cells were pooled to generate 5-cell samples in 10 µL of lysis buffer (SMART-Seq v4 Ultra Low Input RNA Kit for Sequencing, Cat# 634888, Takara Bio, Kusatsu, Japan). RNA sequencing was performed by DNAFORM (Kanagawa, Japan) as previously described[2].

**Dendritic Spine Imaging**

Eight to ten weeks old Arc-Cre$^{ER}$:Ai14 male mice were fed with HFD or RCD for 12 weeks after labeling VMH$^{GE}$ by TRAP experiments. Mice were intracardial perfused with heparinized saline followed by 4 %PFA and brain was fixed in 4%PFA overnight. For assessing effects wnt signaling, mice were sacrificed 3 hours after i.c.v. injection of RSPO1(1ug in 0.5uL) or Dkk1(100ng in 0.5uL). Coronal brains sections contain the VMH were collected with 200um each. Z-stacked sequencing images of dendritic fraction of tdTomato positive neurons in VMH were obtained with Z-steps of 0.5µm by the two-photon excitation laser scanning microscopy (TPLSM) system (A1R-MP+, FN-1, Nikon, Tokyo, Japan) under water-immersion objective lens (Nikon, Apo LWD 25×/1.10 NA). A Ti:Sapphire laser (MaiTai eHP DeepSee, Spectra Physics) was employed as an excitation laser light source at a wavelength of 1000 nm for tdTomato. All fluorescence signals were detected by nondescanned detectors equipped with GaAsP PMTs (non-descanned detector; NDD).

**Western Blotting**

Eight-week-old male mice were maintained on either an RCD or an HFD for 12 weeks. Following euthanasia via $CO_2$ asphyxiation, brains were rapidly extracted, and 1-mm-thick coronal brain sections containing the VMH were prepared using a brain mold. The VMH was microdissected using a surgical blade, immediately snap-frozen in liquid nitrogen, and stored until biochemical analysis. Tissues were homogenized at 4°C in PBS containing 1% Nonidet P-40 (NP-40) and protease inhibitor cocktail (Cat#03969-21, Nacalai Tesque, Kyoto, Japan). Following centrifugation at 15,000 × *g* for 15 min at 4°C, the supernatant was collected and fractionated by SDS-PAGE. Proteins were then transferred onto Immobilon polyvinylidene fluoride (PVDF) membranes (Immobilon; Millipore, MA, USA). The membranes were incubated overnight at 4°C with primary antibodies against total β-catenin (1 µg/mL; sc-7963, Santa Cruz, Dallas, TX) or PSD95 (1:1000; PSD95-Go, Frontier Institute, Hokkaido, Japan), alongside a mouse anti-β-actin antibody (Cell Signaling Technology, MA) as a loading control. Immunoreactive complexes were visualized using horseradish peroxidase (HRP)-

conjugated goat anti-rabbit or anti-mouse IgG secondary antibodies and enhanced chemiluminescence (ECL) reagents (Cytiva, MA, USA).

**Arterial and Venous Catheterization for clamp studies.**
Mice were anesthetized with a mixture of ketamine (100 mg/kg) and xylazine (10 mg/kg). Polyethylene catheters were implanted into the right carotid artery and the jugular vein for blood sampling and infusions, respectively, as previously described[3]. The distal ends of the catheters were tunneled subcutaneously and exteriorized through a small incision at the nape of the neck. Mice were allowed to recover for 3–5 days prior to the clamp experiments. During the recovery period, catheters were flushed daily with heparinized saline to maintain patency.

**Hyperinsulinemic–euglycemic clamp and measurement of 2-[$^{14}$C] deoxy-D-glucose (2DG) uptake.**
Hyperinsulinemic-euglycemic clamps were performed as previously described[4,5]. Experiments were conducted in a freely moving state. A 90-min basal period ($t$ = -90 to 0 min) was followed by a 115-min clamp period ($t$ = 0–115 min). Recombinant RSPO1 protein (1 μg in 0.5 μL) was administered i.c.v. 30 min prior to the basal period ($t$ = -120 min). At the start of the basal period (t = -90 min), a bolus of [3-$^{3}$H] glucose (5 μCi) was injected via the jugular vein, followed by a continuous infusion at 0.05 μCi/min for 90 min. Blood samples were collected at $t$ = -15 and -5 min to determine the basal glucose rate of appearance (Ra). The clamp period was initiated with a continuous insulin infusion (2.5 mU/kg/min). Arterial blood glucose levels were monitored every 5–10 min, and a variable rate of unlabeled "cold" glucose was infused via the jugular vein to maintain euglycemia (110–130 mg/dL). To minimize volume loss, erythrocytes from collected blood samples were resuspended in sterile saline and returned to the animal. To assess tissue-specific glucose uptake, a bolus of 2-[$^{14}$C] DG (10 μCi) was administered at t = 70 min, and blood samples were collected at t = 75, 85, 95, 105, and 115 min. At t = 115 min, mice were euthanized via intravenous thiopental sodium (Nipro, Osaka, Japan), and tissues (soleus muscle, heart, spleen, and hypothalamus) were rapidly harvested and snap-frozen.The whole-body glucose utilization rate (Rd), endogenous glucose production (EGP), and the rates of whole-body glycolysis and glycogen synthesis were calculated as described previously[6].

### QUANTIFICATION AND STATISTICAL ANALYSIS
Statistical analyses were performed using GraphPad Prism 10 software (GraphPad). Comparisons between two groups were analyzed using the two-tailed unpaired Student’s *t*-test. For multiple-group comparisons, one-way or two-way analysis of variance (ANOVA) was used. Repeated-measures ANOVA followed by Sidak’s

multiple comparisons test was applied to analyze data collected over multiple time points. All data are presented as mean ± SEM. A $p$-value $< 0.05$ was considered statistically significant. Representative images were obtained from at least three independent experiments.

#### References

1. Ting, J. T. *et al.* Preparation of Acute Brain Slices Using an Optimized N-Methyl-D-glucamine Protective Recovery Method. *J. Vis. Exp. JoVE* 53825 (2018) doi:10.3791/53825.
2. Imoto, D. *et al.* Refeeding activates neurons in the dorsomedial hypothalamus to inhibit food intake and promote positive valence. *Mol. Metab.* **54**, 101366 (2021).
3. Abe, T. *et al.* Hypothalamic Prostaglandins Facilitate Recovery From Severe Hypoglycemia but Exacerbate Recurrent Hypoglycemia in Mice. *Diabetes* **74**, 2390–2404 (2025).
4. Abe, T. & Toda, C. Hyperglycemic Clamp and Hypoglycemic Clamp in Conscious Mice. *J. Vis. Exp. JoVE* https://doi.org/10.3791/65581 (2024) doi:10.3791/65581.
5. Toda, C. *et al.* UCP2 Regulates Mitochondrial Fission and Ventromedial Nucleus Control of Glucose Responsiveness. *Cell* **164**, 872–883 (2016).
6. Ayala, J. E., Bracy, D. P., McGuinness, O. P. & Wasserman, D. H. Considerations in the design of hyperinsulinemic-euglycemic clamps in the conscious mouse. *Diabetes* **55**, 390–397 (2006).
